# Comprehensive multi-omics study of the molecular perturbations induced by simulated diabetes on coronary artery endothelial cells

**DOI:** 10.1101/2021.01.06.425584

**Authors:** Aldo Moreno-Ulloa, Hilda Carolina Delgado-De la Herrán, Carolina Álvarez-Delgado, Omar Mendoza-Porras, Rommel A. Carballo-Castañeda, Francisco Villarreal

**Affiliations:** MS2 laboratory, Biomedical Innovation Department, Center for Scientific Research and Higher Education of Ensenada (CICESE), Baja California, México; Specialized Laboratory in Metabolomics and Proteomics (MetPro), CICESE, México; Mitochondrial Biology Laboratory, Biomedical Innovation Department, Center for Scientific Research and Higher Education of Ensenada (CICESE), Baja California, México; CSIRO Livestock and Aquaculture, Queensland Bioscience Precinct, 306 Carmody Rd, St Lucia, QLD, Australia; School of Medicine, University of California, San Diego, CA, USA; San Diego VA Healthcare System

**Keywords:** SWATH-Proteomics, Metabolomics, Type 2 Diabetes Mellitus, Endothelial cells, Feature-Based Molecular Networking

## Abstract

Coronary artery endothelial cells (CAEC) exert an important role in the development of cardiovascular disease. Dysfunction of CAEC is associated with cardiovascular disease in subjects with type 2 diabetes mellitus (T2DM). However, comprehensive studies of the effects that a diabetic environment exerts on this cellular type scarce. The present study characterized the molecular perturbations occurring on cultured bovine CAEC subjected to a prolonged diabetic environment (high glucose [HG] and high insulin [HI]). Changes at the metabolite and peptide level were assessed by untargeted metabolomics and chemoinformatics, and the results were integrated with proteomics data using published SWATH-based proteomics on the same *in vitro* model. Our findings were consistent with reports on other endothelial cell types, but also identified novel signatures of DNA/RNA, aminoacid, peptide, and lipid metabolism in cells under a diabetic environment. Manual data inspection revealed disturbances on tryptophan catabolism and biosynthesis of phenylalanine-based, glutathione-based, and proline-based peptide metabolites. Fluorescence microscopy detected an increase in binucleation in cells under treatment that also occurred when human CAEC were used. This multi-omics study identified particular molecular perturbations in an induced diabetic environment that could help unravel the mechanisms underlying the development of cardiovascular disease in subjects with T2DM.

## 1. Introduction

Damage to coronary artery endothelial cells (CAEC) leads to coronary endothelial dysfunction, which is associated with the development of cardiac pathologies in subjects with and without coronary atherosclerosis (1). Subjects with type 2 diabetes mellitus (T2DM) are particularly at increased risk of myocardial infarction (2) and coronary endothelial dysfunction has been implicated in the prognosis (3). A high-glucose (HG) environment—hallmark of T2DM—leads to nitric oxide signaling, cell cycle (4), apoptosis (5), angiogenesis (6), and DNA structure impairment (7). However, given the intrinsic heterogeneity of the endothelium, the molecular perturbations caused by HG vary accordingly with the type of studied endothelial cells (8, 9). For instance, human microvascular endothelial cells showed increased gene expression of endothelial nitric oxide synthase, superoxide dismutase 1, glutathione peroxidase 1, thioredoxin reductase 1 and 2 compared to the regulation observed in human umbilical vein endothelial cells (HUVEC) when cultured in HG for 24 h. Furthermore, the response of endothelial cells to HG is influenced by the duration of exposure (10, 11) as demonstrated in bovine aortic and human microvascular endothelial cells where cell proliferation and apoptosis were higher at <48 h compared to 8 weeks of exposure (10). In another example of time-dependent response, increased apoptosis (derived from DNA fragmentation) and tumor necrosis factor alpha protein levels were reported in human coronary artery endothelial cells (HCAEC) after only 24 h of incubation with HG (5). Hence, the molecular response to HG cannot be generalized among endothelial cell types. Previously we reported impaired mitochondrial function/structure and nitric oxide signaling in HG treated HCAEC for 48 h (12). However, a 72 h study documented an increased in pro-inflammatory cytokines (13) and oxidative stress in HCAEC (14). The long-term (>72 h) effect of HG in CAEC has not been as extensively documented compared to other endothelial cell types. Characterizing the effect of HG on CAEC may allow us to identify key signaling pathways (or specific biomolecules) associated with the development of endothelial dysfunction and cardiac pathologies.

Here, liquid chromatography coupled to mass spectrometry (LC-MS^2^)-based untargeted metabolomics and SWATH-based quantitative proteomics data, as well as bio- and chemo-informatics were used to characterize the molecular perturbations occurring in Bovine Coronary Artery Endothelial Cells (BCAEC) under a prolonged diabetic environment.

## 2. Methods

### 2.1 Chemical and reagents

Recombinant human insulin was purchased from Sigma Aldrich (St. Louis, MO, USA). Antibiotic-antimitotic solution, trypsin-EDTA solution 0.25%, Hank’s Balanced Salt Solution (HBSS) without phenol red, Dulbecco’s Modified Eagle’s Media (DMEM) with glutamine, Fetal Bovine Serum (FBS), Hoechst 33258, Pentahydrate (bis-Benzimide)-FluoroPure^™^, and methanol-free formaldehyde (16% solution) were obtained from Thermo Fisher Scientific (Waltham, MA, USA). Methanol, Acetonitrile, and water were Optima^™^ LC-MS Grade and obtained from Fisher Scientific (Hampton, NH, USA). Ethanol LiChrosolv^®^ Grade was obtained from Merck KGaA (Darmstadt, Germany). Rabbit anti-Von Willebrand factor (vWf) antibody and goat anti-rabbit IgG conjugated to Alexa Fluor 488 were obtained from Abcam (Cambridge, MA, USA).

### 2.2 Cell culture

BCAEC were purchased from Cell applications, Inc. (San Diego, CA, USA) and grown as previously described (15). In brief, cells were grown with DMEM (5.5 mmol/L glucose, supplemented with 10% FBS and 1% antibiotic-antimitotic solution) at 37°C in an incubator with a humidified atmosphere of 5 % CO_2_. Before experiments, cells were switched to DMEM with 1% FBS for 12 h to maintain the cells under a quiescent state. The model to simulate diabetes is described in (15) (**Figure 1**). Endothelial cells were cultured for 12 days to determine the chronic molecular perturbations caused by simulated diabetes and to avoid the early (within 48 h) cell proliferation effects caused by HG (10, 16). In brief, cells were first treated with 100 nmol/L insulin (high-insulin, HI) in normal glucose (NG, 5.5 mmol/L in DMEM) for 3 days (17) and then maintained in high-glucose (HG, 20 mmol/L in DMEM) and constant HI for 9 days. This sequential scheme tried to mimic the pathophysiological conditions that occur in T2DM patients, wherein hyperinsulinemia precedes hyperglycemia (18). Cells were used at passages between 6 to 12. The control group did not receive HI nor HG treatment. For selected experiments (binucleation analysis), HCAEC (55 years old Caucasian male, history of T2DM for >5 years) were purchased from Cell Applications, Inc. and subjected to the same conditions as BCAEC but using MesoEndo Growth Medium (Cell Applications, Inc.) to induce proliferation. For simulated diabetes, HCAEC were treated with HI and HG as with BCAEC but, MesoEndo Growht Medium was used instead. For consistency, the group that underwent simulated diabetes (HG + HI) will be referred to as the “experimental group”. All experiments were carried out in triplicate.

**Figure 1.**
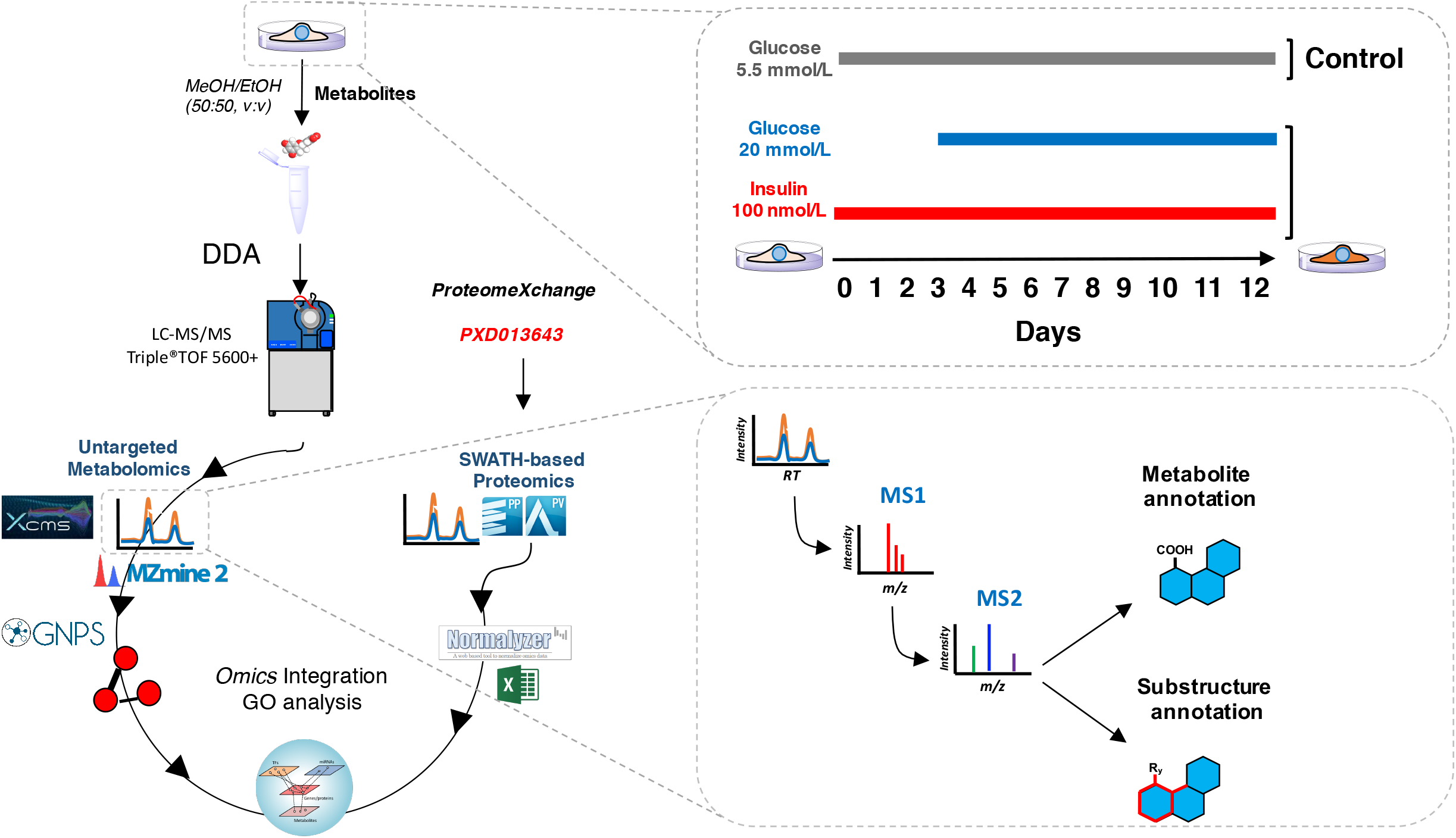
Illustration of the methodology followed in this study.

### 2.3 Immunofluorescence

As previously described (15), 100,000 cells per well were seeded onto 12-well plates (Corning^®^ CellBIND^®^) and exposed to simulated diabetes. Thereafter, BCAEC and HCAEC were washed with PBS to remove dead cells and debris. Cells were fixed, permeabilized, and blocked as described before (19). Cells were then incubated with a polyclonal antibody against the vWf (1:400, 3% BSA in PBS) overnight at 4°C and thereafter washed 3x with PBS. Alexa Fluor 488-labeled anti-rabbit (1:400 in PBS) was then used as a secondary antibody for 1 h at RT and washed 3x with PBS. As a negative control, cells were incubated only with secondary antibody to assess for non-specific binding. Cell nuclei were stained with Hoechst 33258 (2 μg/ml in HBSS) for 30 min and washed 3x with PBS. Fluorescent images were taken in at least three random fields per condition using an EVOS^®^ FLoid^®^ Cell Imaging Station with a fixed 20x air objective. Image analysis was performed through ImageJ software (version 2.0.0).

### 2.4 Metabolite extraction

Cells were seeded at 300,000 cells per well in 6-well plates (Corning^®^ CellBIND^®^) and treated as above. After HG and HI conditions, metabolites were extracted following a published protocol for adherent cells with some modifications (20) (**Figure 1**). In brief, after washing the cells 3 x with PBS, 500 μL of a cold mixture of methanol: ethanol (50:50, v:v) were added to each well, covered with aluminum foil, and incubated at −80°C for 4 h. Cells were then scrapped using a lifter (Fisher Scientific, Hampton, NH, USA), and the supernatant was transferred to Eppendorf tubes before centrifugation for 10 min at 14,000 rpm at 4°C. The supernatant was transferred to another tube and dried down by SpeedVac^™^ System (Thermo Fisher Scientific, Waltham, MA, USA). Samples were reconstituted in water/acetonitrile 95:5 v/v with 0.1% formic, centrifuged at 14,000 rpm for 10 min at 4°C. The particle free supernatant was recovered for further LC-MS^2^ analysis.

### 2.5 LC-MS^2^ data acquisition for metabolomics

Metabolites were loaded into an Eksigent nanoLC^®^ 400 system (AB Sciex, Foster City, CA, USA) with a HALO Phenyl-Hexyl column (0.5 x 50 mm, 2.7 μm, 90 Å pore size, Eksigent AB Sciex, Foster City, CA, USA) for data acquisition using the LC-MS parameters previously described with some modifications (21). In brief, the separation of metabolites was performed using gradient elution with 0.1% formic acid in water (A) and 0.1% formic acid in ACN (B) as mobile phases at a constant flow rate of 5 μL/min. The gradient started with 5% B for 1 min followed by a stepped increase to 100%, B over 26 min and held constant for 4 min. Solvent composition was returned to 5% B for 0.1 min. Column re-equilibration was carried out with 5% mobile phase B for 4 minutes. Potential carryover was minimized with a blank run (1 μL buffer A) between sample experimental samples. The eluate from the LC was delivered directly to the TurboV source of a TripleTOF 5600+ mass spectrometer (AB Sciex, Foster City, CA, USA) using electrospray ionization (ESI) under positive mode. ESI source conditions were set as follows: IonSpray Voltage Floating, 5500 V; Source temperature, 350°C; Curtain gas, 20 psi; Ion source gases 1 and 2 were set to 40 and 45 psi; Declustering potential, 100 V. Data was acquired using information-dependent acquisition (IDA) with high sensitivity mode selected, automatically switching between full-scan MS and MS/MS. The accumulation time for TOF MS was 0.25 s/spectra over the *m/z* range 100-1500 Da and for MS/MS scan was 0.05 s/spectra over the *m/z* 50-1500 Da. The IDA settings were as follows charge state +1 to +2, intensity 125 cps, exclude isotopes within 6 Da, mass tolerance 50 mDa, and a maximum number of candidate ions 20. Under IDA settings, the ‘‘exclude former target ions’’ was set as 15 s after two occurrences and ‘‘dynamic background subtract’’ was selected. Manufacturer rolling collision energy (CE) option was used based on the size and charge of the precursor ion using formula CE=*m/z* x 0.0575 + 9. The instrument was automatically calibrated by the batch mode using appropriate positive TOF MS and MS/MS calibration solutions before sample injection and after injection of two samples (<3.5 working hours) to ensure a mass accuracy of <5 ppm for both MS and MS/MS data. Instrument performance was monitored during data acquisition by including QC samples (pooled samples of equal volume) every 4 experimental samples. Data acquisition of experimental samples was also randomized.

### 2.6 Metabolomics data processing

Mass detection, chromatogram building and deconvolution, isotopic assignment, feature alignment, and gap-filling (to detect features missed during the initial alignment) from LC-MS^2^ datasets was performed using XCMS (https://xcmsonline.scripps.edu) (22) and MZmine (23) software. The XCMS pipeline was used for normalization of feature area and statistical analysis. To identify or annotate the metabolites at the chemical structure and class level, the MS^2^-containing features extracted with MZmine were further analyzed using the Global Natural Products Social Molecular Networking (GNPS) (24), Network Annotation Propagation (NAP) (25) and MS2LDA (26) *in silico* annotation tools, and Classyfire automated chemical classification (27), as previously described (21) with some modifications. The confidences of such annotations are level 2 (probable structure by library spectrum match) and level 3 (tentative candidates) in agreement with the Metabolomics Standards Initiative (MSI) classification (28). Molecular networking, NAP, and Classyfire outputs were integrated using the MolNetEnhancer workflow (29). Molecular networks were visualized using Cytoscape version 3.8.2 (30). In addition, chemical substructures (co-occurring fragments and neutral losses referred to as “mass2motifs” [M2M]) were recognized using the MS2LDA web pipeline (http://www.ms2lda.org) to further annotate metabolites (level 3, MSI). The detailed processing parameters for XCMS and MZmine pipelines are found in the supporting information.

### 2.7 Peptidomics data processing

For peptide identification, raw .wiff and .wiff.scan files (same files used for MZmine and XCMS) from the experimental and control groups were analyzed separately using ProteinPilot software version 4.2 (Ab Sciex, Foster City, CA, USA) with the Paragon algorithm. MS^1^ and MS^2^ data were searched against the *Bos taurus* SwissProt sequence database (6006 reviewed proteins+common protein contaminants, February 2019 release). The parameters input was: sample type, identification; digestion, none; Cys alkylation, none; instrument, TripleTOF 5600; special factors, none; species, *Bos taurus;* ID focus, biological modifications, and amino acid substitutions; search effort, thorough ID. False discovery rate analysis was also performed. All peptides were exported and those with a >90% confidence were linked to the corresponding feature extracted by the XCMS algorithm using their accurate mass and retention time information. For peptide quantification, we employed the normalized feature abundances (MS^1^ level) generated by XCMS. A significance threshold of *p<0.05* (Welch’s t test) was utilized.

### 2.8 Proteomics data reprocessing

The SWATH-based proteomics data (identifier PXD013643), hosted in ProteomeXchange consortium via PRIDE (31), was reanalyzed with some modifications. The parameters used to build the spectral library remained the same (15), while the parameter for peptides per protein was set to 100 in the software SWATH^®^ Acquisition MicroApp 2.0 in PeakView^®^ version 1.2 (AB Sciex, Foster City, CA, USA). The obtained protein peak areas were exported to Markerview^™^ version 1.3 (AB Sciex, Foster City, CA, USA) for further data refinement, including assignment of IDs to files and removal of reversed and common contaminants. Peak areas were exported in a .tsv file, and normalized with NormalyzerDE online version 1.3.4 (32). The NormalyzerDE pipeline comprises 8 different normalization methods (Log2, variance stabilizing normalization, total intensity, median, mean, quantile, CycLoess, and robust linear regression). The results of qualitative (MA plots, scatter plots, box plots, density plots) and quantitative (pooled intragroup coefficient of variation [PCV], median absolute deviation [PMAD], estimate of variance [PEV]) parameters were compared between the normalization methods to select the most appropriate.

### 2.9 Bioinformatic analysis of proteomics data

Proteins that passed the significance threshold were first converted to their corresponding Entrez Gene (GeneID) using https://www.uniprot.org/uploadlists/ and then transformed to their human equivalents using the ortholog conversion feature in https://biodbnet-abcc.ncifcrf.gov/db/dbOrtho.php. Bioinformatic analysis was done on OmicsNet website platform (https://www.omicsnet.ca/) (33, 34). First, a protein-protein interaction (PPI) molecular network (first-order network containing query or seeds molecules and their immediate interacting partners) using STRING PPI database was built (35) and then pathway enrichment analysis was performed using the built-in REACTOME and the Kyoto Encyclopedia of Genes and Genomes (KEGG) databases. To visualize modules (functional units) contained in the molecular network the WalkTrap algorithm (within OmicsNet platform) was employed. Hypergeometric test was used to compute *p-values*.

### 2.10 Integrative analysis of proteomics and metabolomics data

The molecular interactions between the proteins and metabolites differentially abundant between HG + HI and NG were determined in OmicsNet (32, 33). The lists of proteins (EntrezGene ID) and metabolites (HMDB ID) were loaded to build a composite network using protein-protein (STRING database selected) and metabolite-protein (KEGG database selected) interaction types. The primary network relied on the metabolite input. Pathway enrichment analysis was performed using the built-in REACTOME and KEGG databases. Hypergeometric test was used to compute *p-values*.

### 2.11 Statistical analysis

All experiments were performed in triplicate. Based on the accuracy (determination of real fold-changes) of SWATH-based quantification (36), proteins with a fold change ≥ 1.2 or ≤ 1/1.2 and a *p-value* <0.05 (Welch’s t-test) were considered as differentially abundant between NG and HG + HI conditions. For the metabolomics data, features with a fold change ≥ 1.3 or ≤ 1/1.3 and a *p-value* <0.05 (Welch’s t-test) were considered as differentially abundant. We did not apply multiple-test corrections to calculate adjusted p-values, because this process could obscure proteins or metabolites with real changes (true-positives) (37). Instead, the analysis was focused on top-enriched signaling pathways (adjusted *p-value* <0.01) that allowed us to determine a set of interacting proteins and metabolites with relevant biological information and contributes in reducing false positives. For multivariate statistical analysis and heatmap visualization, Metaboanalyst 4.0 (https://www.metaboanalyst.ca) was utilized. Principal component analysis (PCA) was used to assess for sample clustering behavior and inter-group variation. No scaling was used for PCA and heatmap analysis. Software PRISM 6.0 (GraphPad Software, San Diego, CA) was used for the creation of volcano plots and column graphs.

### 2.12 Data availability

The raw datasets supporting the metabolomics results are available in the GNPS/MassIVE public repository (38) under the accession number MSV000084307. The specific parameters of the tools employed for metabolite annotation are available on the following links: for classical molecular networking, https://gnps.ucsd.edu/ProteoSAFe/status.jsp?task=604b3d077e00430a9bc288eebf154b9b; for FBMN https://gnps.ucsd.edu/ProteoSAFe/status.jsp?task=5e2839037969442e868d9df21309d561; for NAP, https://proteomics2.ucsd.edu/ProteoSAFe/status.jsp?task=96cda48c0df64d3398a8f9088907afb5; for MS2LDA, http://ms2lda.org/basicviz/summary/1197/ (need to log-in as a registered or guest user); for MolNetEnhancer, https://gnps.ucsd.edu/ProteoSAFe/status.jsp?task=de80b9c765e042ffab7767a3101054fd. The quantitative results generated using the XCMS platform can be accessed after logging into the following link https://xcmsonline.scripps.edu and searching for the job number 1395724. SWATH data is accessible on the ProteomeXchange with dataset identifier PXD013643.

## 3. Results

### Untargeted metabolomics

Overall 5571 features or potential metabolites were detected using XCMS and MZmine, wherein 957 (~18%) features were commonly identified in both platforms (**Figure 2A**). Based on the relative quantification using XCMS, 140 and 82 features were detected with reduced and increased abundances respectively in the experimental group compared to the control group (**Figure 2B**). The effects of HG and HI in the experimental group are observed by PCA analysis wherein the experimental samples clustered away from the control group (**Figure 2C**). The consistency of the LC-MS equipment is apparent by the clustering of the QC samples (**Figure 2C**). Further, the heatmap visualization of the top 100-modulated metabolites exhibited the different distribution patterns among groups (**Figure 2D**). Using the GNPS platform for automatic metabolite annotation, 106 compounds (excluding duplicates and contaminants) were putatively annotated with a level 2 confidence annotation (MS^2^ spectral match) (**Table S1**) in agreeance with the MSI classification (28). Some metabolites identified by the GNPS platform could not be quantified because they were not detected by the XCMS algorithm during feature area normalization and quantification. Moreover, GNPS Molecular Networking aligned the MS^2^-containing features (n=1,013) based on their structural similarity, creating 118 independent networks or clusters with at least two connected nodes (**Figure 3A**). The use of MolNetEnhancer workflow allowed to putatively identify chemical classes (level 3, MSI) for 56 of the 118 independent networks. The top-10 most abundant annotated chemical classes and associated metabolites are shown in **Figure 3A.** Three-clusters from the network were further analyzed because they contained annotated metabolites by spectral matching, which facilitates the annotation of other cluster’s nodes. Cluster 1 revealed two metabolites linked to the organonitrogen compounds class with reduced abundance in the experimental group (**Figure 3B**). Library spectral match (level 2, MSI) suggest PC(16:0/18:1(9Z)) and PC(18:0/18:2(9Z,12Z)) as putative candidates, which was supported by MS2LDA phosphocholine-substructure recognition (**Figure 3C**). In cluster 2, glutathione-based metabolites (MSI level 3) were detected through fragments *m/z* 308.0925, 233.0575, 179.0475, and 162.0225 retrieved by the M2M_453 substructure and associated with glutathione structure using mzCloud *in silico* predictions (**Figure 4A**). The precursor ion at *m/z* 713.1472 and glutathione (annotated at level 2, MSI) were detected with increased abundance in the experimental group. MS2LDA visualization, at the M2M level, correlated with the GNPS molecular networking clustering (**Figure 4B**). In cluster 3, various phenylalanine-based metabolites were putatively annotated aided by MS2LDA substructure recognition (**Figure 4C and 4D**). Within this cluster, glutamyl-phenylalanine (annotated at level 2, MSI) and the precursor ions at *m/z* 297.1802 and 487.1548 presented with increased abundance in the experimental vs. control group. On the other hand, various aminoacids were annotated (level 2, MSI) by GNPS spectral matching and manual inspection of data (**Table S2**). Threonine, valine, proline, leucine, serine, glutamic acid, methionine, and tyrosine presented increased abundance (fold change range 1.3-1.7, p<0.05) in the experimental vs. control group. Particularly, metabolites linked to the catabolism of tryptophan via the serotonin and kynurenine pathway (39) were annotated (level 2, MSI), including melatonin, acetyl serotonin, and kynurenine (**Table S1**). However, only kynurenine was significantly elevated in the experimental group. The full list of annotated metabolites, differential abundances and another relevant feature information is shown in **Table S2**.

**Figure 2.**
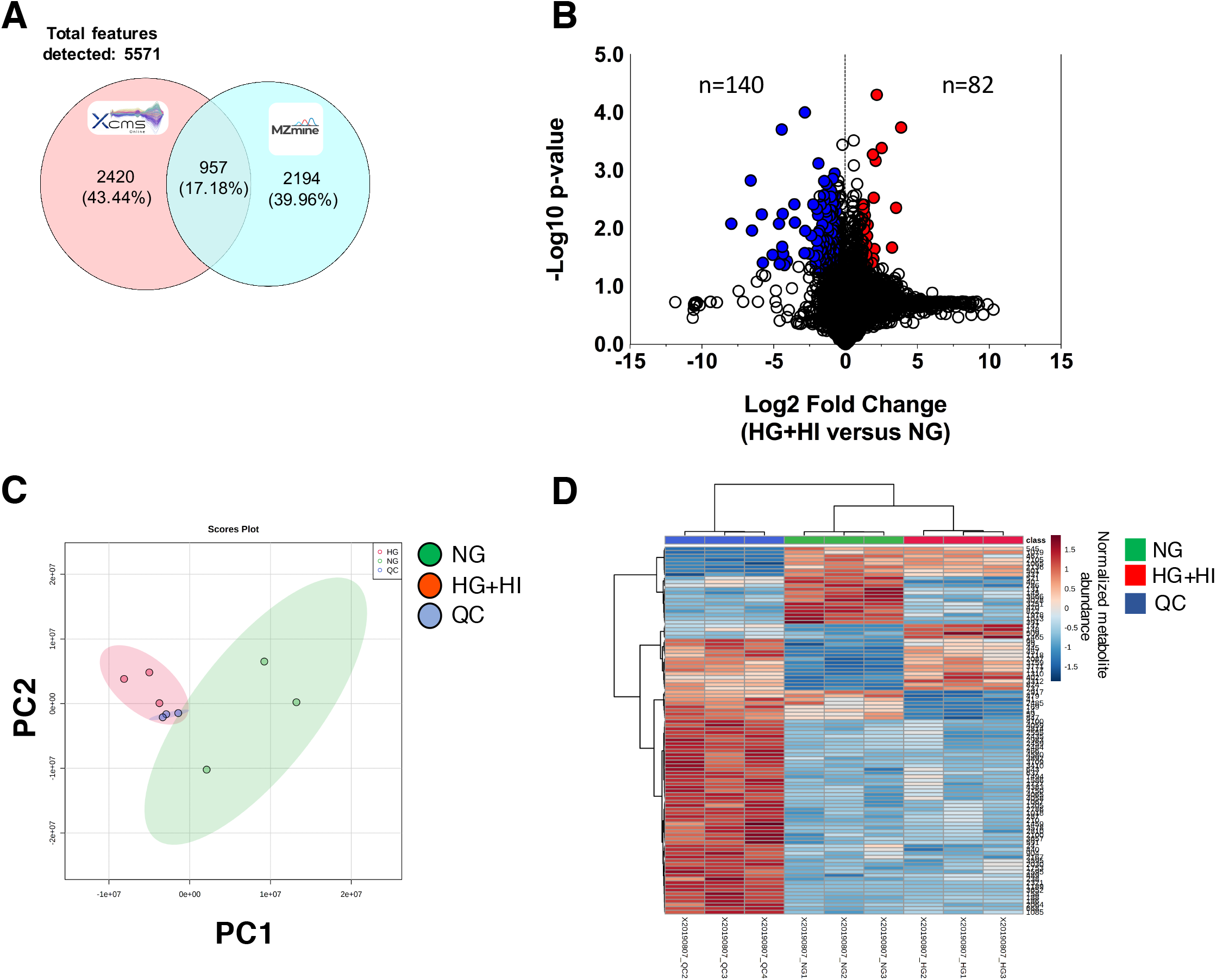
Simulated diabetes induced changes in the metabolome of bovine coronary artery endothelial cells (BCAEC). (A) Venn diagram of features identified among MZmine and XCMS software (0.01 Da and 1 min retention time, thresholds) on LC-MS^2^ datasets. (B) Volcano plot of all quantified metabolites displaying differences in relative abundance (> +/−30% change, <0.05 *p-value* cut-offs) between BCAEC cultured in control (NG) media and simulated diabetes (HG+ HI) for twelve days. Values (dots) represent the HG+HI/NG ratio for all metabolites. Red and blue dots denote downregulated and upregulated metabolites in the HG + HI group vs. NG group, respectively. (C) Principal Component Analysis (PCA) of LC-MS^2^ datasets. Data was log transformed without scaling. Shade areas depict the 95% confidence intervals. (C) HeatMap of the top 100 metabolites ranked by t-test. Abbreviations: NG, normal glucose; HG, high glucose; HI, high insulin; QC, quality control.

**Figure 3.**
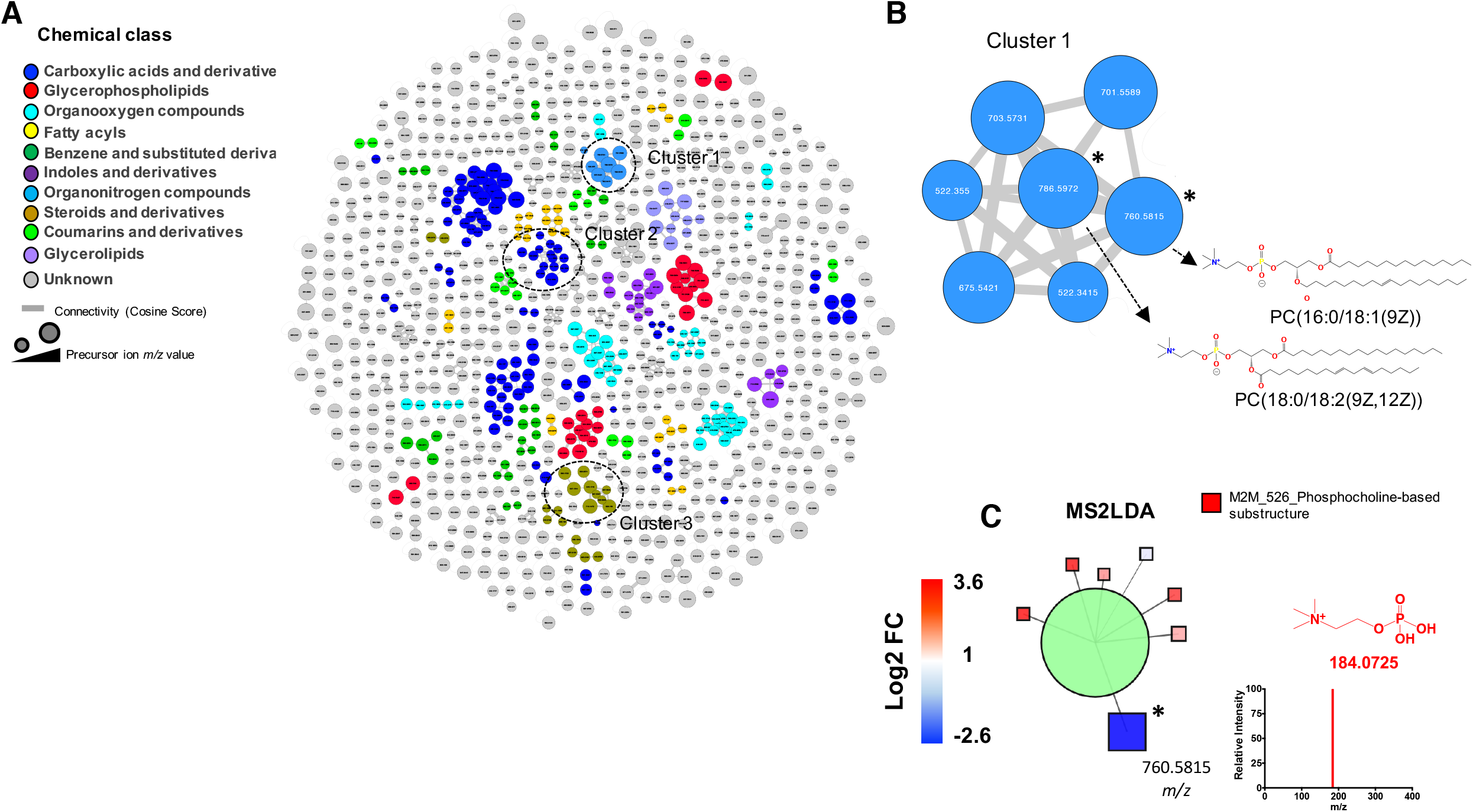
Bovine coronary artery endothelial cells (BCAEC) metabolite molecular network. (A) Molecular classes (according to Classyfire) of the metabolome identified by the MolNetEnhancer workflow and visualized by Cytoscape version 3.8.2. Each node represents a unique feature and the color of the node denotes the associated chemical class. The thickness of the edge (connectivity) indicates the MS^2^ similarity (Cosine score) among features. The *m/z* value of the feature is shown inside the node and is proportional to the size of the node. Three selected clusters or connected features as relevant are shown. (B) Inset of cluster 1 denoting the presence of phosphocholine (PC)-containing lipids. Significant differential abundant features among simulated diabetes (HG+HI) and control (NG) groups are indicated with an asterisk (*p-value* <0.05). (C) Characterization of features in (B) aided by substructure recognition by MSLDA software using MS^1^ visualization in www.ms2lda.org. Fragment at *m/z* 184.0725 linked to a PC head group by mzCloud *in silico* prediction (www.mzCloud.org). Abbreviations: M2M, mass2motif; FC, fold change; NG, normal glucose; HG, high glucose; HI, high insulin. Chemical structures were drawn by ChemDraw Professional version 16.0.1.4.

**Figure 4.**
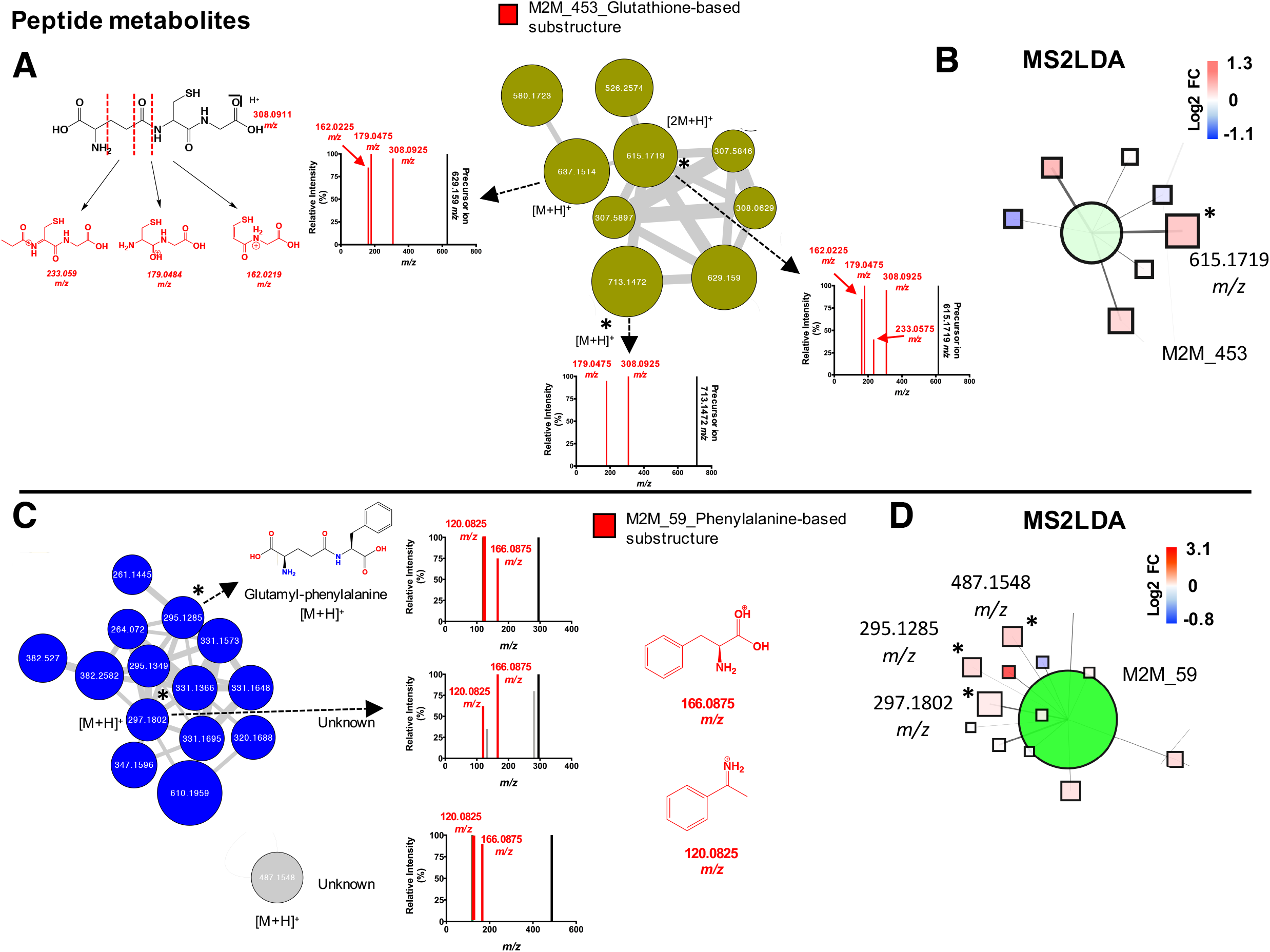
Peptide metabolites modulated by simulated diabetes in bovine coronary artery endothelial cells (BCAEC). (A) Cluster 2 retrieved from the main molecular network linked to glutathione and derivatives. The fragments of mass-2-motif (M2M)_453 colored in red are characteristic of a glutathione core and the fragments are shown in red. (B) Features associated with M2M_453 using MS^1^ visualization in www.ms2lda.org. (C) Cluster 3 retrieved from the main molecular network linked to phenylalanine-based metabolites. A singular node at *m/z* 487.1548 is also shown. The fragments of M2M_59 colored in red are characteristic of a phenylalanine core (Heuristic and Quantum Chemical predictions by www.mzCloud.org). (D) Features associated with M2M_59 using MS^1^ visualization in www.ms2lda.org. In GNPS’s clusters (A and C), the node’s color denotes the chemical class assigned to the cluster. The thickness of the edge (connectivity) indicates the cosine score (MS^2^ similarity). The *m/z* value of the feature is shown inside the node and is proportional to the size of the node. Significant differential abundant features among simulated diabetes (HG+HI) and control (NG) groups are indicated with an asterisk (*p-value* <0.05). In MS2LDA’s nodes (B and D), the green node represents the M2M and squares indicate individual features. Edges represent connections to M2M. Significant differential abundant features among groups are indicated with an asterisk (*p-value* <0.05). Abbreviations: M2M, mass2motif; FC, fold change; NG, normal glucose; HG, high glucose; HI, high insulin. Chemical structures were drawn by ChemDraw Professional version 16.0.1.4.

### Peptidomics

Experimental and control datasets were analyzed separately to identify the peptides and their biological modifications. The complete list of peptides identified by ProteinPilot between the experimental and control groups are described in **Table S3.** Proline oxidation was the most frequent biological modification detected in the experimental group datasets. We identified 8 and 12 peptides with a confidence of >90% in the control and experimental group, respectively. Differential abundance of 2 proline-rich peptides was observed in the experimental group compared to the control group. An additional tripeptide was manually annotated with a LPP sequence (**Table S4**).

### Proteomics

The re-analysis of the SWATH data (PXD013643 dataset) facilitated the identification of 952 quantifiable proteins (717 proteins with at least 2 unique peptides, 1% false discovery rate) and no missing values among technical and biological replicates (**Table S5**). Sample datasets were normalized using 8 different methods to select the most appropriate based on quantitative and qualitative parameters on our dataset. Quantile normalization produced a better qualitative and quantitative profile and was selected to further process our data (**Figure S1**). PCA analysis of normalized data denoted a clear separation of the groups suggesting overall differences in their proteomes (**Figure 5A**). Differential abundance analysis revealed 32 and 33 proteins with increased and decreased abundance in the experimental group (**Figure 5B**). Further, the heatmap visualization of the top 50-modulated proteins exhibited the different distribution patterns among the experimental and control groups (**Figure 5C**). To obtain a molecular insight we performed a functional enrichment analysis using a network-based approach. First, we created a composite network comprising PPI between the modulated proteins by simulated diabetes (seed proteins) and their immediate interacting partners (highest confidence >0.9) retrieved from STRING Database (incorporated in OmicsNet platform). The principal network using the up-modulated proteins consisted of 461 proteins, 709 edges and 18 seed proteins (nodes with blue shadow) and is illustrated in **Figure 5D**. Eight modules or clusters were generated, that may represent relevant complexes or functional units (40). The 5 most significant (adjusted p-value <0.05) REACTOME and KEGG pathways on the global network are shown in **Table 1**. Two modules contained multiple seed proteins and were linked to DNA/RNA and protein metabolism pathways using the WalkTrap algorithm (**Figure 5D**). On the other hand, the principal network using the down-modulated proteins consisted of 488 proteins, 513 edges and 18 seed proteins identified eleven modules wherein one module (with 2 seed proteins) indicated associations with mitochondrial function pathways (**Figure 5E**).

**Figure 5.**
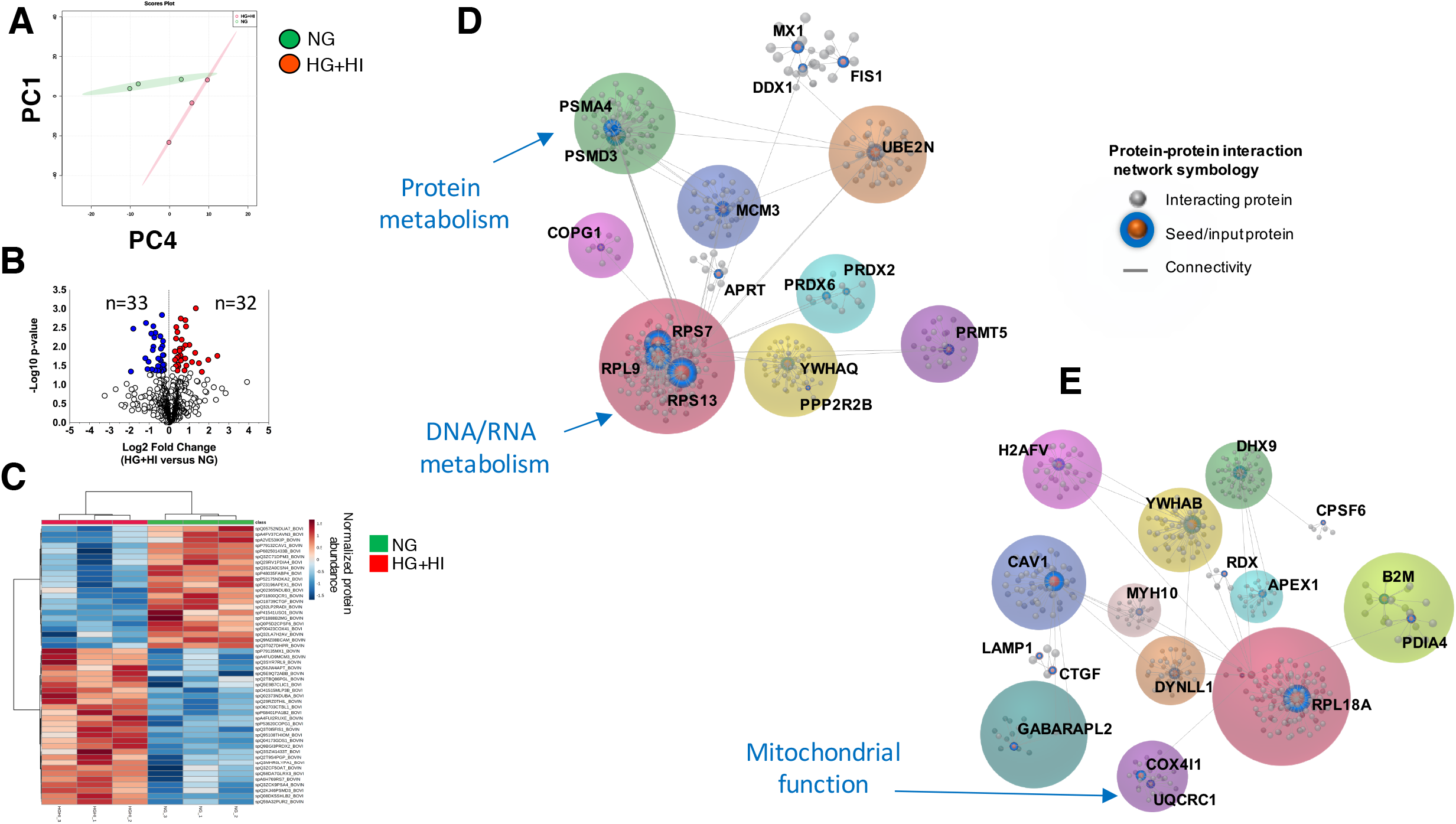
Simulated diabetes induced changes in the proteome of bovine coronary artery endothelial cells (BCAEC). (A) Principal Component Analysis (PCA) of LC-SWATH-MS^2^ datasets. Data was log transformed without scaling. Shade areas depict the 95% confidence intervals. No scaling was used. (B) Volcano plot of all quantified proteins (Quantile normalization) displaying differences in relative abundance (> +/−20% change, <0.05 *p-value* cut-offs) between BCAEC cultured in control (NG) media and simulated diabetes (HG+ HI) for twelve days. Values (dots) represent the HG+HI/NG ratio for all proteins. Red and blue dots denote downregulated and upregulated proteins in the HG + HI group vs. NG group, respectively. (C) HeatMap of the top 50 metabolites ranked by t-test. Protein-Protein interactome (>0.9 confidence) using the list of proteins with increased abundance (D) and reduced abundance (E) in the HG + HI group. Colored circles denote modules or clusters which may represent relevant complexes or functional units. The input proteins are illustrated with a blue shade and the gene ID is also shown. The most representative pathway (containing more input proteins) for all modules is indicated in blue letters. Abbreviations: NG, normal glucose; HG, high glucose; HI, high insulin.

**Table 1.**
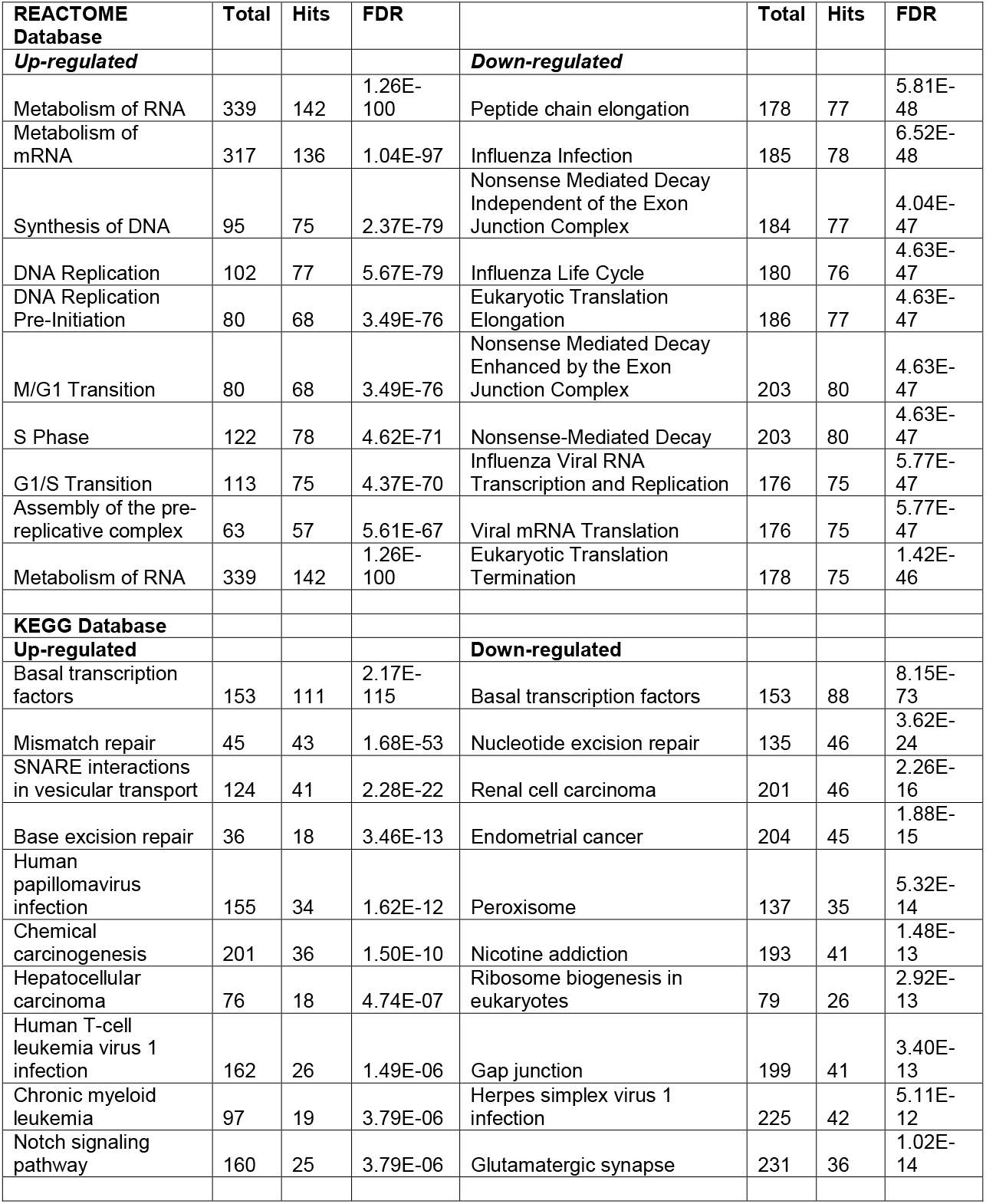
Pathway enrichment analysis of up-regulated and down-regulated proteins in HG+HI group.

### Integration of Metabolomics and Proteomics

The signaling pathways perturbed by simulated diabetes were identified by a composite network of interacting metabolites and proteins using OmicsNet built-in databases. **Figure 6** illustrates the composite bi-layered metabolite-PPI network using the up-modulated molecules (under simulated diabetes) comprised of 9 metabolites (seed metabolites), 177 edges, and 166 proteins (5 seed proteins). The 10 top-most enriched signaling pathways identified in the composite network are shown in **Table 2**. The two principal modules highlighted by the WalkTrap algorithm were linked to glutathione and amino acid metabolism. We noted a smaller interaction between Acyl-protein thioesterase 1 (LYPLA1) and a phosphatidylcholine metabolite when simultaneously analyzing up- and down-modulated proteins and metabolites. No significant composite network was identified using the down-modulated proteins and metabolites.

**Figure 6.**
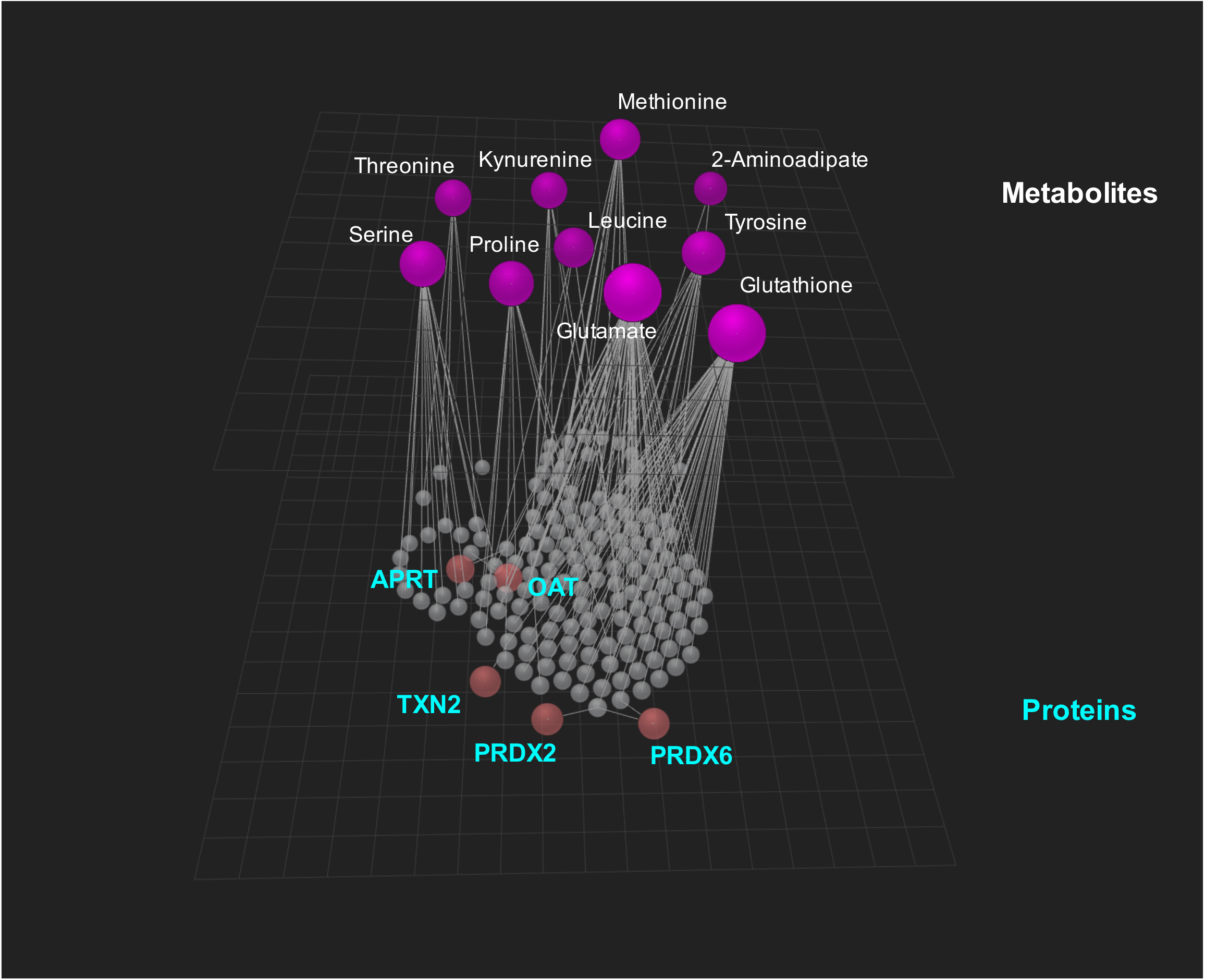
3D Integrative network of the proteomic and metabolomic perturbations caused by simulated diabetes in bovine coronary artery endothelial cells (BCAEC). Composite protein-metabolite network created by OmicsNet using the up-regulated proteins (red nodes) and metabolites (magenta nodes) in the HG + HI group (simulated diabetes). Interacting proteins (<0.9 confidence) were retrieved from STRING Database and are shown as gray nodes. Abbreviations: NG, normal glucose; HG, high glucose; HI, high insulin.

**Table 2.**
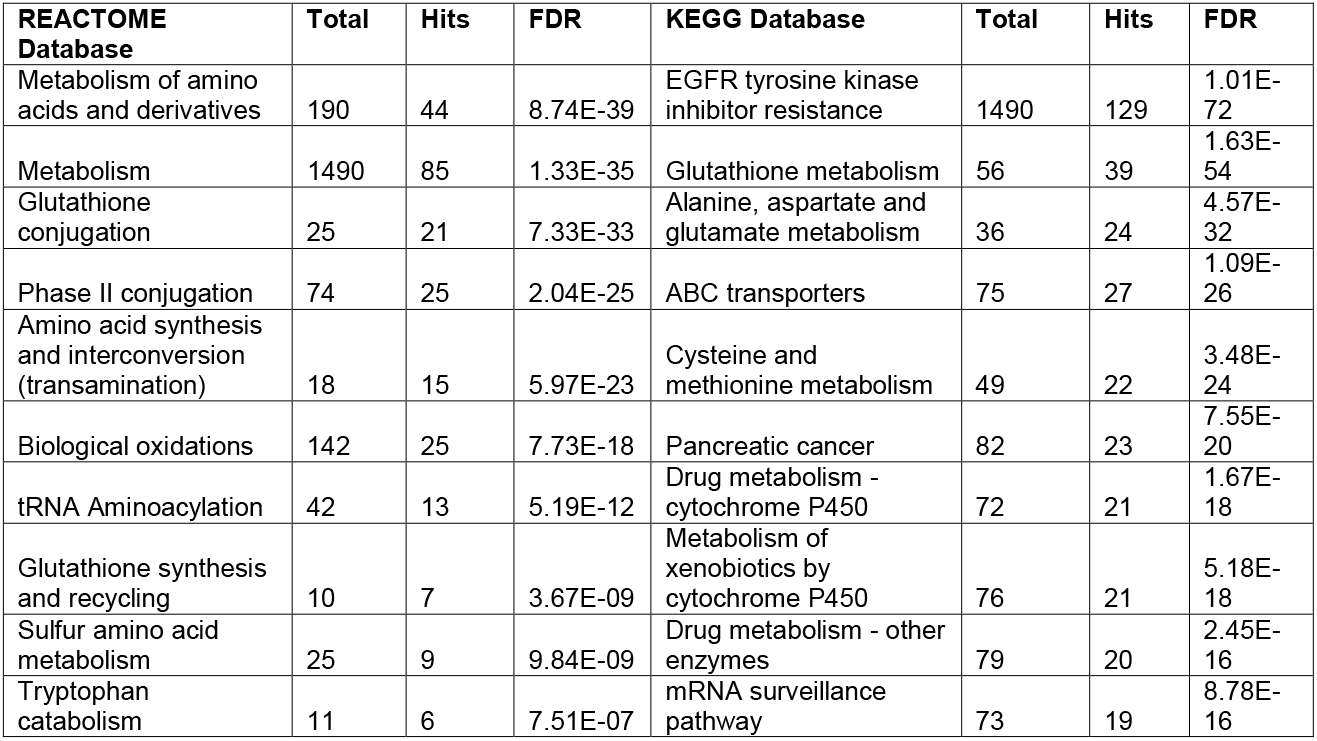
Integrative pathway enrichment analysis of up-regulated proteins and metabolites in HG+HI group.

### Cellular morphology

To better understand the effects that simulated diabetes exerts on endothelial cells the changes on cellular structure endpoints were evaluated. The endothelial nuclei morphology in the BCAEC control and experimental groups were evaluated using fluorescent-staining and image analysis. We also evaluated the presence of vWF (marker of endothelial cells) in BCAEC and HCAEC, to reveal the cellular boundary and to demonstrate their endothelial phenotype (41). We noted an increase in the percentage of binucleated BCAEC in the experimental group compared to the control group (top panel **Figure 7A and 7B**). A similar result with larger nuclei, was observed when using HCAEC as a human *in vitro* model (bottom panel **Figure 7A and 7B**). Finally, as expected, we observed a typical intracellular localization of vWF and a 100% positivity in endothelial cells.

**Figure 7.**
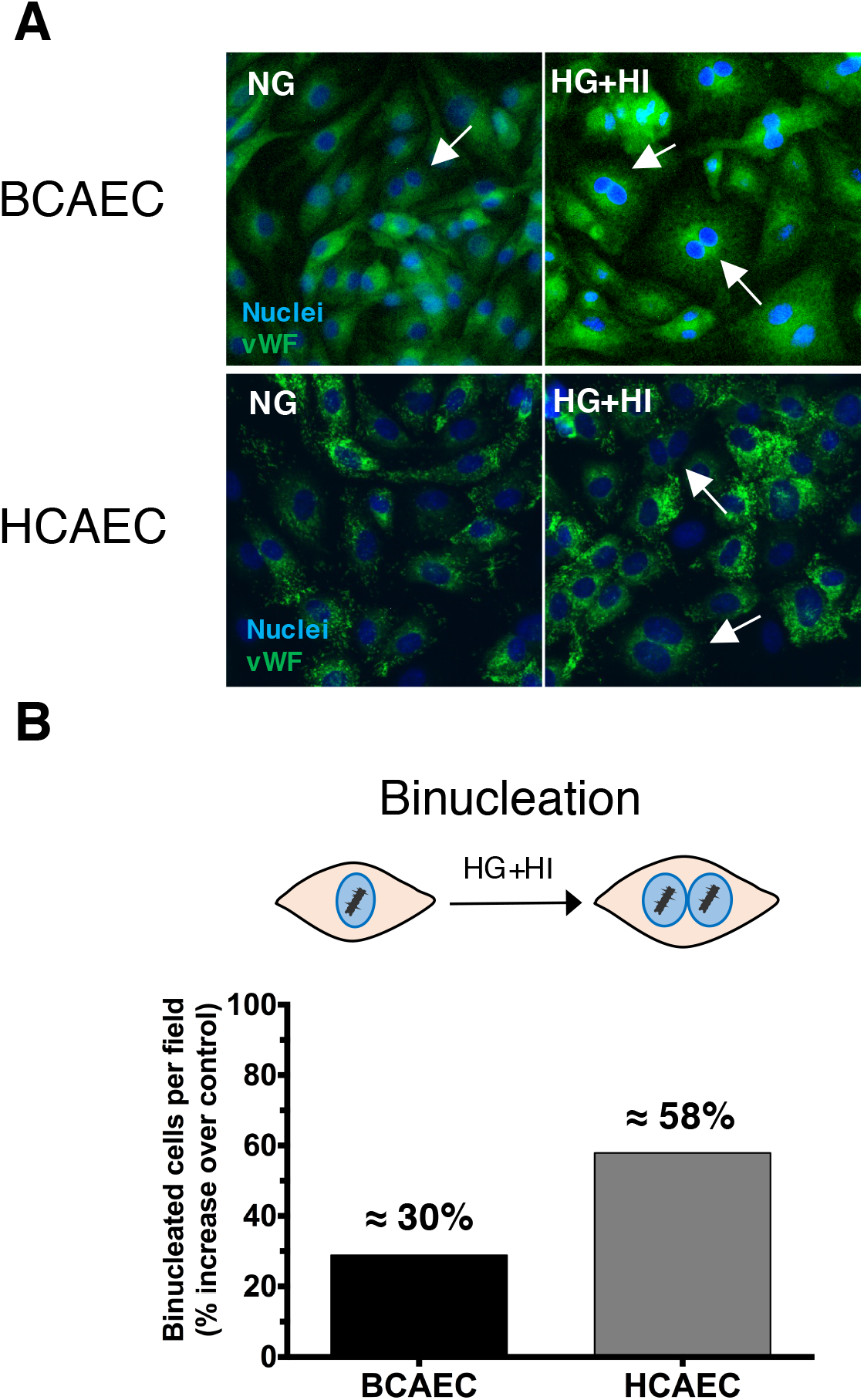
Increased cellular binucleation by simulated diabetes in bovine coronary artery endothelial cells (BCAEC) and human coronary artery endothelial cells (HCAEC). (A) Representative immunofluorescence micrographs showing the localization of the von-Willebrand factor (vWf, 1:400, 3% BSA in PBS) in fixed and permeabilized cells. The nuclei were stained using the dye Hoechst 33258 (2 μg/ml in HBSS). White arrows indicate binucleated cells. (B) Quantification of binucleated cells in HCAEC and BCAEC under simulated diabetes (HG+HI) vs. control (NG) group. Fluorescence images were taken in at least three random fields per condition using an EVOS^®^ FLoid^®^ Cell Imaging Station with a fixed 20x air objective. Image analysis was performed by ImageJ software (version 2.0.0). Abbreviations: NG, normal glucose; HG, high glucose; HI, high insulin.

## 4. Discussion

This study investigated the molecular perturbations occurring in coronary endothelium cells subjected to prolonged simulated diabetes that facilitated the identification of signaling pathways and specific molecules that could be associated with the development of cardiovascular disease. To achieve this, we employed a MS-based multi-omics approach coupled to fluorescence microscopy to detect structural changes. Endothelial cells cover the inner surface of blood vessels and are distributed across the body. Their functions include: acting as a mechanical barrier between the circulating blood and adjacent tissues as well as modulating multiple functions in distinct organs (42). These regulatory functions vary according to localization and vascular bed-origin (43). HG blood levels are detrimental to endothelial cells function in T2DM leading to coronary endothelial dysfunction and development of CVD (44, 45). The molecular effects of HG on endothelial cells have been previously characterized (4, 6, 7, 10, 11); nevertheless, the endothelial cell types used in these studies are not intrinsically involved in CVD. The present study used an *in vitro* model involving endothelial cells that modulate the heart function, CAEC (46). Our model not only used HG (20 mmol/L) to simulate diabetes (4, 6, 7, 10, 11) but first induced insulin resistance to mimic the pathophysiological conditions that occur in T2DM wherein hyperinsulinemia precedes hyperglycemia (18). Diabetes was simulated for up to 12 days to mimic chronic HG exposure and to prevent measuring cell proliferation known to occur in early HG (10, 16). Despite a lack of apparent increase in cell proliferation in the experimental group compared to control group after twelve days, an increase in overall protein abundance was detected by Bradford assay (data not shown) and inferred from total ion chromatogram (TIC) of MS (**Figure S1A**). We suggest that protein synthesis is increased as a consequence of the higher presence of bi-nucleated CAEC (with increased DNA/RNA metabolism) under HG + HI compared to that in the control cohort (**Figure 7A and 7B**). Previous studies have shown reduced endothelial cell proliferation (mostly in HUVEC) after long-term (7-14 days) HG exposure (4, 11, 47–53), accompanied by an increase in protein synthesis (53). This MS-based methodological pipeline that included appropriate controls during data acquisition (QC) and processing (e.g., normalization, filtering, annotation, dereplication, etc.), allowed the identification of global changes in the metabolome of CAEC under HG + HI. Specifically, increased abundance of valine, leucine, tyrosine, serine, leucine, proline, methionine, and glutamic acid in cells under HG conditions was observed; and this is consistent with reports on human aortic endothelial cells (54). Notably, several clinical studies have established a direct relationship between prevalence/incidence of T2DM and increased levels of valine, leucine and tyrosine in serum and plasma (55–59). Our results support the role of CAEC in contributing to the elevated pool of amino acids seen in circulation under a HG environment. We speculate that increased levels of these amino acids could result from either increased production or reduced degradation as suggested in endothelial cells (immortalized cell line, EA.hy 926) that transition from a glycolytic metabolism towards lipid and amino acid oxidation when challenged by HG (60). Furthermore, evidence of increased tryptophan catabolism was identified through the kynurenine pathway. In this regard, a non-significant decrease of ~ 40% in the abundance of tryptophan was detected. However, a significant increase of ~ 450% in kynurenine (tryptophan’s main metabolite) (61) between the HG + HI group and NG group was also observed, which is a key finding as elevated plasma levels of kynurenine are known to increase CVD risk (62, 63). This novel finding contributes to expanding the understanding of amino acid metabolism in endothelial cells under simulated diabetes. Acetyl serotonin and melatonin which are components of the serotonin pathway that degrades tryptophan (64) were also detected with only minor abundancy increases (20-30%) in the HG + HI group compared to control. Differences in glutathione (cysteine-glutamic acid-glycine, tripeptide) metabolism in CAEC were also found, suggesting an increased response to oxidative stress (65). In line with this observation, previous research reported a glutathione-dependent reaction to ambient HG in artery-derived endothelial cells (66, 67) but the same could not be observed in vein-derived endothelial cells (68, 69). This emphasizes the different responses to HG among endothelial phenotypes. Here, novel evidence is provided of the up-regulation of glutathione-based metabolites. The composite protein network suggested an increase in glutathione metabolism supported by elevated levels of oxidized glutathione and, one of its synthetic precursors, glutamic acid. At the protein level, peroxiredoxin (PRDX2 and PRDX6) and thioredoxin (TXN2, mitochondrial) showed increased abundances in the experimental group, which are part of the cells natural enzymatic defense against oxidative stress (70). The substructure analysis of metabolomics data facilitated identifying glutamic acid- and phenylalanine-based metabolites, presumably di- or tri-peptides, including the annotated metabolite glutamyl-phenylalanine. Furthermore, the CAEC peptidome analysis suggested an increase in proline-containing peptides. This type of peptide is of particular interest because of their resistance to non-specific proteolytic degradation, body distribution and remarkable biological effects (71–74). Yet, the precise function of such phenylalanine-, glutamine-, and proline-based peptides remains to be characterized in CAEC. We can only speculate that they are the result of a compensatory mechanism to reduce glucose cellular damage. Also, increased protein abundance of core and regulatory subunits from the proteasome complex (PSMA4 and PSMD3) was found in cells under simulated diabetes. This suggests an increased protein degradation and subsequent peptide formation in response to HG. Metabolomic profiling also revealed changes in the lipidome of CAEC challenged with HG + HI, wherein a reduction in phosphatidylcholine (PC) lipids and subsequent increase in phosphocholine were noted. Changes in the phospholipidomic profile of bovine aortic endothelial cells treated with HG for 24 h has also been reported in a lipidome study (75). Here, proteomics and metabolomics data were manually integrated and this allowed to determine critical roles for PAFAH1B2 and LYPLA1 in mediating the degradation of PC lipids (**Figure 8**). PAFAH1B2 was found to be up-regulated in this study and it is known to be associated with inflammation and higher levels of lysoPC (76). As a result, PAFAH1B2 could increase the pool of lysoPC lipids, further exacerbating inflammation in the cardiovascular system (77). On the other hand, LYPLA1 has a lysophospholipase activity that can hydrolyze a range of lysophospholipids, including LysoPC, thereby generating a fatty acid and glycerophosphocholine as products (78). Increased levels of phosphocholine (~ 460%) were detected in HG treated cells compared to control, that could be associated with the degradation of LysoPC lipids. It should be noted that the use of pathways databases such as KEGG and REACTOME possess some limitations when dealing with lipid metabolites because its chemical diversity is not well annotated/defined within the databases. For example, KEGG provides a chemical class identifier instead of individual identity to lipids, constricting their biological importance (79). Thus, based on our manual inspection of the metabolomics-proteomics data and in line with the evidence, we suggest that simulated diabetes evokes inflammation on BCAEC and that PAFAH1B2 and LYPLA1 play a role in modulating such process.

**Figure 8.**
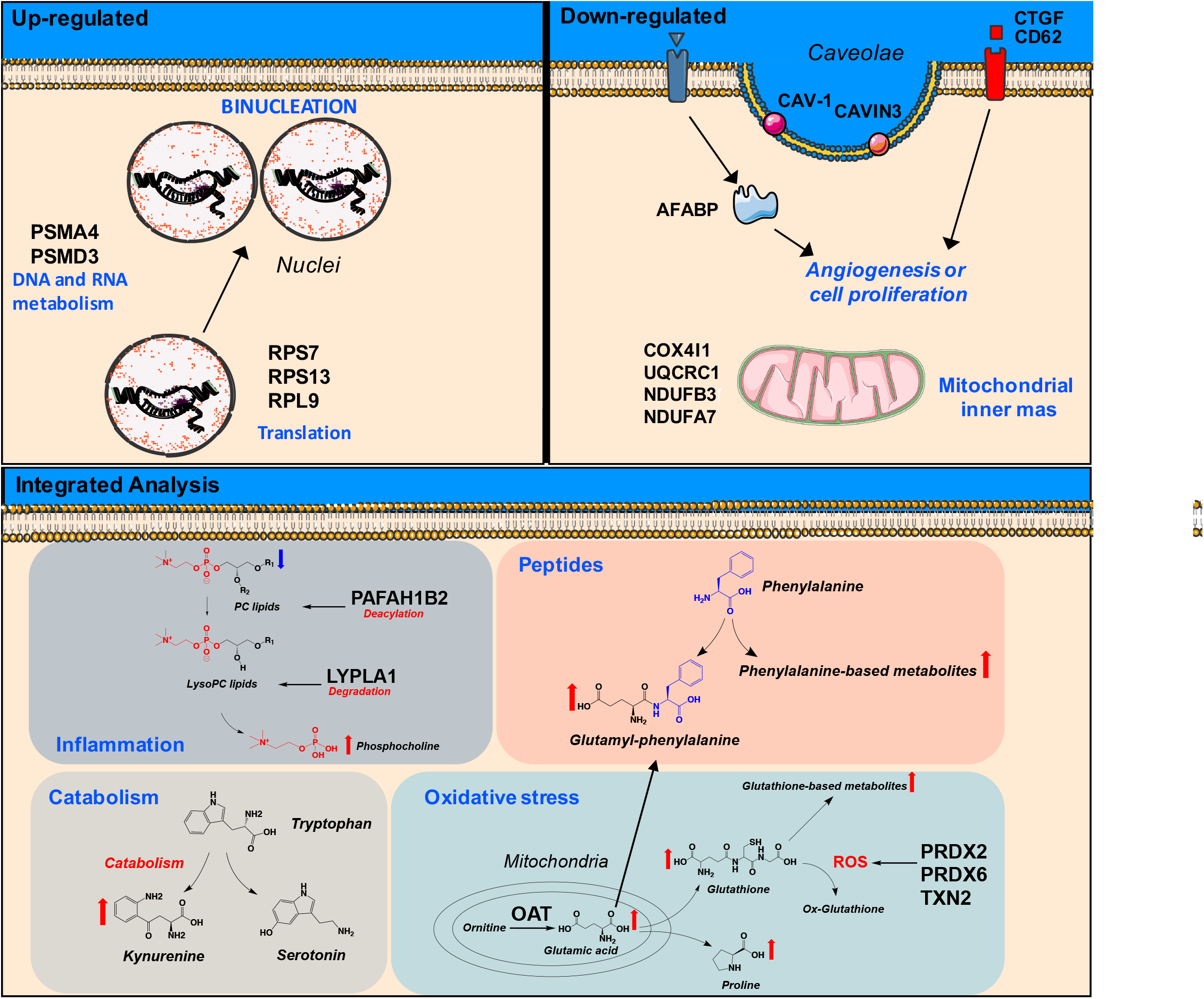
Summary illustration of study findings. Cellular structures were created using Servier Medical Art templates, which are licensed under a Creative Commons Attribution 3.0 Unported License; https://smart.servier.com. Chemical structures were drawn by ChemDraw Professional version 16.0.1.4.

Previously, we reported the multinucleation of CAEC cultured under simulated diabetes (15). This type of cell possesses ≥2 nuclei. Here, we replicated our previous findings of increased binucleation in BCAEC. The same outcome was obtained when using HCAEC as a human *in vitro* model (**Figure 7A and 7B**), validating the binucleation process in other CAEC. After refinement of LC-MS^2^ data and bioinformatics re-processing of published SWATH-based datasets of BCAEC under simulated diabetes (15), molecular signatures and pathways that could be linked to the binucleation process were found (**Figure 8**). For instance, we noted an increased abundance of proteins, under simulated diabetes, with reported nuclei localization and linked to DNA metabolism, including ribosomal proteins RPS7, RPS13, and RPL9 (80). Further, we observed an increased abundance of proteasome proteins, PSMA4 and PSMD3, which are linked to protein metabolism (81). Hence, we infer that the CAEC binucleation occurs as a compensatory mechanism to increase the cell capacity to metabolize the excess of ambient glucose by increasing the cell metabolic machinery (transcription/translation processes). Although an increase in cell proliferation could boost a coordinated increase of ribosomal and proteasome proteins, we do not believe this is the case here, as mentioned before. After 4-5 days of simulated diabetes, cells occupied 100% of the well’s plate surface, thereby impeding to harbor more cells because endothelial cells grow as a monolayer. This is consistent with findings stating that when endothelial cells become highly confluent, they stop growing due to cell-cell contact, even in the presence of growth factors (82). In support of this, up-stream (CTGF and CD62) (83, 84) (**Table S5**) and down-stream proteins (FABP4) (85) (**Table S5**) involved in angiogenesis and proliferation were down-regulated by simulated diabetes. Importantly, there is evidence (not in endothelial cells) of cellular processes contributing to the stimulation of cellular binucleation without increases in cell proliferation, including cellular enhancement of antimicrobial defenses (86), senescence (87), and malignancy (88). Various mechanisms have been linked to the binucleation process, such as cytokinesis failure, cellular fusion, mitotic slippage, and endoreduplication (89). The elucidation of the exact molecular mechanisms leading to the binucleation process of CAEC is beyond the scope of our study.

In conclusion, this study applied an integrated multi-omics and bioinformatics/chemoinformatics approach to characterize the molecular perturbations that simulated diabetes exerts on CAEC. We confirmed several independent studies that reported alterations at protein and metabolite levels in endothelial cells of different sources than coronary vessels. Metabolomics, identified alterations in amino acid, peptide, and phospholipid metabolism. Notably, the chemoinformatic analysis identified unreported alterations of phenylalanine-, glutathione-, and proline-based peptides on coronary endothelium under simulated diabetes. Proteomics provided evidence of reduced mitochondrial mass and angiogenesis. The integration of proteomics and metabolomics identified increased glutamic acid metabolism and suggested that the antioxidant enzymes are involved in protecting the cells from oxidative stress. Fluorescence microscopy reported the appearance of non-proliferative binucleated CAEC cells as a mean to metabolize the excess of ambient glucose. Overall, our study improved the understanding of the molecular disturbances caused by simulated diabetes that could mediate CAEC dysfunction and may be relevant in the context of CVD in subjects with T2DM.

## 5. Acknowledgements

This work was derived in part from the Thesis Project of H.C.D.H. at the Posgrado en Ciencias de la Vida, CICESE. We thank Alan G. Hernández-Melgar for his invaluable technical assistance with the NormalyzerDE software.

## 6. Funding

Part of this work was supported by CICESE (Grant No. 685109 to AMU and Internal Project No. 685-110 from CAD), NIH R01 DK98717 (to FV), and VA Merit-I01 BX3230 (to FV).

## 7. Conflict of interest

Dr. Villarreal is a co-founder and stockholder of Cardero Therapeutics, Inc.

## 8. Author contributions

A.M.U. contributed to the study conception and design, data acquisition, formal analysis, methodology, project administration, and funding acquisition. H.C.D.H., L.D.M, and R.A.C.C. contributed to the data acquisition, formal analysis and interpretation of some experiments. C.A.D., and F.V. contributed to funding acquisition and resources. O.M.P contributed to data interpretation and critical revision of manuscript. All authors contributed to the drafting, revising, and approval of the final version of the manuscript.

## Supporting information

**Table S1.** List of all the putatively annotated metabolites by MS^2^ spectral matching against GNPS public spectral libraries.

**Table S2.** List of putatively annotated (MS^2^ spectral matching) metabolites modulated by simulated diabetes.

**Table S3.** List of all detected peptides by ProteinPilot Software using the metabolomics datasets.

**Table S4** Putative annotated proline-peptides altered by simulated diabetes in Bovine Coronary Artery Endothelial Cells by ProteinPilot Software and manual inspection.

**Table S5.** List of the detected peptides and proteins in all conditions for SWATH-based quantification.

**Figure S1.** Proteomics data normalization results using NormalyzerDE. (A) Total intensity of raw data before normalization. (B) Quantitative parameters of normalization algorithms (pooled intragroup coefficient of variation [PCV], median absolute deviation [PMAD], estimate of variance [PEV]). Qualitative parameters of normalization algorithms; (C) Box plots (D) MA plots, and (E) Density plots.

**Figure S2. Cellular confluence in control and experimental group.** Representative micrographs of Bovine Coronary Artery Endothelial Cells (BCAEC) cultured for 9 days with 5.5 mmol/L glucose (control group) and 20 mmol/L glucose+100 nmol/L insulin (simulated diabetes or experimental group). Images were taken using an EVOS^®^ FLoid^®^ Cell Imaging Station with a fixed 20x air objective. Abbreviations: NG, normal glucose; HG, high glucose; HI, high insulin.

## References

1. Halcox, J. P.; Schenke, W. H.; Zalos, G.; Mincemoyer, R.; Prasad, A.; Waclawiw, M. A.; Nour, K. R.; Quyyumi, A. A., Prognostic value of coronary vascular endothelial dysfunction. Circulation 2002, 106, (6), 653–8.

2. Lind, M.; Wedel, H.; Rosengren, A., Excess Mortality among Persons with Type 2 Diabetes. N Engl J Med 2016, 374, (8), 788–9.

3. Gutierrez, E.; Flammer, A. J.; Lerman, L. O.; Elizaga, J.; Lerman, A.; Fernandez-Aviles, F., Endothelial dysfunction over the course of coronary artery disease. Eur Heart J 2013, 34, (41), 3175–81.

4. Lorenzi, M.; Cagliero, E.; Toledo, S., Glucose toxicity for human endothelial cells in culture. Delayed replication, disturbed cell cycle, and accelerated death. Diabetes 1985, 34, (7), 621–7.

5. Kageyama, S.; Yokoo, H.; Tomita, K.; Kageyama-Yahara, N.; Uchimido, R.; Matsuda, N.; Yamamoto, S.; Hattori, Y., High glucose-induced apoptosis in human coronary artery endothelial cells involves up-regulation of death receptors. Cardiovasc Diabetol 2011, 10, 73.

6. Dubois, S.; Madec, A. M.; Mesnier, A.; Armanet, M.; Chikh, K.; Berney, T.; Thivolet, C., Glucose inhibits angiogenesis of isolated human pancreatic islets. J Mol Endocrinol 2010, 45, (2), 99–105.

7. Lorenzi, M.; Montisano, D. F.; Toledo, S.; Barrieux, A., High glucose induces DNA damage in cultured human endothelial cells. J Clin Invest 1986, 77, (1), 322–5.

8. Patel, H.; Chen, J.; Das, K. C.; Kavdia, M., Hyperglycemia induces differential change in oxidative stress at gene expression and functional levels in HUVEC and HMVEC. Cardiovasc Diabetol 2013, 12, 142.

9. Pala, L.; Pezzatini, A.; Dicembrini, I.; Ciani, S.; Gelmini, S.; Vannelli, B. G.; Cresci, B.; Mannucci, E.; Rotella, C. M., Different modulation of dipeptidyl peptidase-4 activity between microvascular and macrovascular human endothelial cells. Acta Diabetol 2012, 49 Suppl 1, S59–63.

10. Esposito, C.; Fasoli, G.; Plati, A. R.; Bellotti, N.; Conte, M. M.; Cornacchia, F.; Foschi, A.; Mazzullo, T.; Semeraro, L.; Dal Canton, A., Long-term exposure to high glucose up-regulates VCAM-induced endothelial cell adhesiveness to PBMC. Kidney Int 2001, 59, (5), 1842–9.

11. Baumgartner-Parzer, S. M.; Wagner, L.; Pettermann, M.; Grillari, J.; Gessl, A.; Waldhausl, W., High-glucose--triggered apoptosis in cultured endothelial cells. Diabetes 1995, 44, (11), 1323–7.

12. Ramirez-Sanchez, I.; Rodriguez, A.; Moreno-Ulloa, A.; Ceballos, G.; Villarreal, F., (-)-Epicatechin-induced recovery of mitochondria from simulated diabetes: Potential role of endothelial nitric oxide synthase. Diab Vasc Dis Res 2016, 13, (3), 201–10.

13. Liu, T.; Gong, J.; Chen, Y.; Jiang, S., Periodic vs constant high glucose in inducing pro-inflammatory cytokine expression in human coronary artery endothelial cells. Inflamm Res 2013, 62, (7), 697–701.

14. Liu, T. S.; Pei, Y. H.; Peng, Y. P.; Chen, J.; Jiang, S. S.; Gong, J. B., Oscillating high glucose enhances oxidative stress and apoptosis in human coronary artery endothelial cells. J Endocrinol Invest 2014, 37, (7), 645–51.

15. Hilda Carolina Delgado De la Herrán, L. D.-M., Carolina Álvarez-Delgado, Francisco Villarreal, Aldo Moreno-Ulloa, Formation of multinucleated variant endothelial cells with altered mitochondrial function in cultured coronary endothelium under simulated diabetes. bioRxiv 2019.

16. Li, X. X.; Liu, Y. M.; Li, Y. J.; Xie, N.; Yan, Y. F.; Chi, Y. L.; Zhou, L.; Xie, S. Y.; Wang, P. Y., High glucose concentration induces endothelial cell proliferation by regulating cyclin-D2-related miR-98. J Cell Mol Med 2016, 20, (6), 1159–69.

17. Madonna, R.; De Caterina, R., Prolonged exposure to high insulin impairs the endothelial PI3-kinase/Akt/nitric oxide signalling. Thromb Haemost 2009, 101, (2), 345–50.

18. Zaccardi, F.; Webb, D. R.; Yates, T.; Davies, M. J., Pathophysiology of type 1 and type 2 diabetes mellitus: a 90-year perspective. Postgrad Med J 2016, 92, (1084), 63–9.

19. Moreno-Ulloa, A.; Miranda-Cervantes, A.; Licea-Navarro, A.; Mansour, C.; Beltran-Partida, E.; Donis-Maturano, L.; Delgado De la Herran, H. C.; Villarreal, F.; Alvarez-Delgado, C., (-)-Epicatechin stimulates mitochondrial biogenesis and cell growth in C2C12 myotubes via the G-protein coupled estrogen receptor. Eur J Pharmacol 2018, 822, 95–107.

20. Kirkwood, J. S.; Maier, C.; Stevens, J. F., Simultaneous, untargeted metabolic profiling of polar and nonpolar metabolites by LC-Q-TOF mass spectrometry. Curr Protoc Toxicol 2013, Chapter 4, Unit 4 39.

21. Moreno-Ulloa, A.; Sicairos Diaz, V.; Tejeda-Mora, J. A.; Macias Contreras, M. I.; Castillo, F. D.; Guerrero, A.; Gonzalez Sanchez, R.; Mendoza-Porras, O.; Vazquez Duhalt, R.; Licea-Navarro, A., Chemical Profiling Provides Insights into the Metabolic Machinery of Hydrocarbon-Degrading Deep-Sea Microbes. mSystems 2020, 5, (6).

22. Gowda, H.; Ivanisevic, J.; Johnson, C. H.; Kurczy, M. E.; Benton, H. P.; Rinehart, D.; Nguyen, T.; Ray, J.; Kuehl, J.; Arevalo, B.; Westenskow, P. D.; Wang, J.; Arkin, A. P.; Deutschbauer, A. M.; Patti, G. J.; Siuzdak, G., Interactive XCMS Online: simplifying advanced metabolomic data processing and subsequent statistical analyses. Anal Chem 2014, 86, (14), 6931–9.

23. Pluskal, T.; Castillo, S.; Villar-Briones, A.; Oresic, M., MZmine 2: modular framework for processing, visualizing, and analyzing mass spectrometry-based molecular profile data. BMC Bioinformatics 2010, 11, 395.

24. Aron, A. T.; Gentry, E. C.; McPhail, K. L.; Nothias, L. F.; Nothias-Esposito, M.; Bouslimani, A.; Petras, D.; Gauglitz, J. M.; Sikora, N.; Vargas, F.; van der Hooft, J. J. J.; Ernst, M.; Kang, K. B.; Aceves, C. M.; Caraballo-Rodriguez, A. M.; Koester, I.; Weldon, K. C.; Bertrand, S.; Roullier, C.; Sun, K.; Tehan, R. M.; Boya, P. C.; Christian, M. H.; Gutierrez, M.; Ulloa, A. M.; Tejeda Mora, J. A.; Mojica-Flores, R.; Lakey-Beitia, J.; Vasquez-Chaves, V.; Zhang, Y.; Calderon, A. I.; Tayler, N.; Keyzers, R. A.; Tugizimana, F.; Ndlovu, N.; Aksenov, A. A.; Jarmusch, A. K.; Schmid, R.; Truman, A. W.; Bandeira, N.; Wang, M.; Dorrestein, P. C., Reproducible molecular networking of untargeted mass spectrometry data using GNPS. Nat Protoc 2020.

25. da Silva, R. R.; Wang, M.; Nothias, L. F.; van der Hooft, J. J. J.; Caraballo-Rodriguez, A. M.; Fox, E.; Balunas, M. J.; Klassen, J. L.; Lopes, N. P.; Dorrestein, P. C., Propagating annotations of molecular networks using in silico fragmentation. PLoS Comput Biol 2018, 14, (4), e1006089.

26. van der Hooft, J. J.; Wandy, J.; Barrett, M. P.; Burgess, K. E.; Rogers, S., Topic modeling for untargeted substructure exploration in metabolomics. Proc Natl Acad Sci U S A 2016, 113, (48), 13738–13743.

27. Djoumbou Feunang, Y.; Eisner, R.; Knox, C.; Chepelev, L.; Hastings, J.; Owen, G.; Fahy, E.; Steinbeck, C.; Subramanian, S.; Bolton, E.; Greiner, R.; Wishart, D. S., ClassyFire: automated chemical classification with a comprehensive, computable taxonomy. J Cheminform 2016, 8, 61.

28. Schymanski, E. L.; Jeon, J.; Gulde, R.; Fenner, K.; Ruff, M.; Singer, H. P.; Hollender, J., Identifying small molecules via high resolution mass spectrometry: communicating confidence. Environ Sci Technol 2014, 48, (4), 2097–8.

29. Ernst, M.; Kang, K. B.; Caraballo-Rodriguez, A. M.; Nothias, L. F.; Wandy, J.; Chen, C.; Wang, M.; Rogers, S.; Medema, M. H.; Dorrestein, P. C.; van der Hooft, J. J. J., MolNetEnhancer: Enhanced Molecular Networks by Integrating Metabolome Mining and Annotation Tools. Metabolites 2019, 9, (7).

30. Shannon, P.; Markiel, A.; Ozier, O.; Baliga, N. S.; Wang, J. T.; Ramage, D.; Amin, N.; Schwikowski, B.; Ideker, T., Cytoscape: a software environment for integrated models of biomolecular interaction networks. Genome Res 2003, 13, (11), 2498–504.

31. Perez-Riverol, Y.; Csordas, A.; Bai, J.; Bernal-Llinares, M.; Hewapathirana, S.; Kundu, D. J.; Inuganti, A.; Griss, J.; Mayer, G.; Eisenacher, M.; Perez, E.; Uszkoreit, J.; Pfeuffer, J.; Sachsenberg, T.; Yilmaz, S.; Tiwary, S.; Cox, J.; Audain, E.; Walzer, M.; Jarnuczak, A. F.; Ternent, T.; Brazma, A.; Vizcaino, J. A., The PRIDE database and related tools and resources in 2019: improving support for quantification data. Nucleic Acids Res 2019, 47, (D1), D442–D450.

32. Willforss, J.; Chawade, A.; Levander, F., NormalyzerDE: Online Tool for Improved Normalization of Omics Expression Data and High-Sensitivity Differential Expression Analysis. J Proteome Res 2019, 18, (2), 732–740.

33. Zhou, G.; Xia, J., Using OmicsNet for Network Integration and 3D Visualization. Curr Protoc Bioinformatics 2019, 65, (1), e69.

34. Zhou, G.; Xia, J., OmicsNet: a web-based tool for creation and visual analysis of biological networks in 3D space. Nucleic Acids Res 2018, 46, (W1), W514–W522.

35. Szklarczyk, D.; Franceschini, A.; Wyder, S.; Forslund, K.; Heller, D.; Huerta-Cepas, J.; Simonovic, M.; Roth, A.; Santos, A.; Tsafou, K. P.; Kuhn, M.; Bork, P.; Jensen, L. J.; von Mering, C., STRING v10: protein-protein interaction networks, integrated over the tree of life. Nucleic Acids Res 2015, 43, (Database issue), D447–52.

36. Muntel, J.; Kirkpatrick, J.; Bruderer, R.; Huang, T.; Vitek, O.; Ori, A.; Reiter, L., Comparison of Protein Quantification in a Complex Background by DIA and TMT Workflows with Fixed Instrument Time. J Proteome Res 2019, 18, (3), 1340–1351.

37. Pascovici, D.; Handler, D. C.; Wu, J. X.; Haynes, P. A., Multiple testing corrections in quantitative proteomics: A useful but blunt tool. Proteomics 2016, 16, (18), 2448–53.

38. Wang, M.; Carver, J. J.; Phelan, V. V.; Sanchez, L. M.; Garg, N.; Peng, Y.; Nguyen, D. D.; Watrous, J.; Kapono, C. A.; Luzzatto-Knaan, T.; Porto, C.; Bouslimani, A.; Melnik, A. V.; Meehan, M. J.; Liu, W. T.; Crusemann, M.; Boudreau, P. D.; Esquenazi, E.; Sandoval-Calderon, M.; Kersten, R. D.; Pace, L. A.; Quinn, R. A.; Duncan, K. R.; Hsu, C. C.; Floros, D. J.; Gavilan, R. G.; Kleigrewe, K.; Northen, T.; Dutton, R. J.; Parrot, D.; Carlson, E. E.; Aigle, B.; Michelsen, C. F.; Jelsbak, L.; Sohlenkamp, C.; Pevzner, P.; Edlund, A.; McLean, J.; Piel, J.; Murphy, B. T.; Gerwick, L.; Liaw, C. C.; Yang, Y. L.; Humpf, H. U.; Maansson, M.; Keyzers, R. A.; Sims, A. C.; Johnson, A. R.; Sidebottom, A. M.; Sedio, B. E.; Klitgaard, A.; Larson, C. B.; P, C. A. B.; Torres-Mendoza, D.; Gonzalez, D. J.; Silva, D. B.; Marques, L. M.; Demarque, D. P.; Pociute, E.; O’Neill, E. C.; Briand, E.; Helfrich, E. J. N.; Granatosky, E. A.; Glukhov, E.; Ryffel, F.; Houson, H.; Mohimani, H.; Kharbush, J. J.; Zeng, Y.; Vorholt, J. A.; Kurita, K. L.; Charusanti, P.; McPhail, K. L.; Nielsen, K. F.; Vuong, L.; Elfeki, M.; Traxler, M. F.; Engene, N.; Koyama, N.; Vining, O. B.; Baric, R.; Silva, R. R.; Mascuch, S. J.; Tomasi, S.; Jenkins, S.; Macherla, V.; Hoffman, T.; Agarwal, V.; Williams, P. G.; Dai, J.; Neupane, R.; Gurr, J.; Rodriguez, A. M. C.; Lamsa, A.; Zhang, C.; Dorrestein, K.; Duggan, B. M.; Almaliti, J.; Allard, P. M.; Phapale, P.; Nothias, L. F.; Alexandrov, T.; Litaudon, M.; Wolfender, J. L.; Kyle, J. E.; Metz, T. O.; Peryea, T.; Nguyen, D. T.; VanLeer, D.; Shinn, P.; Jadhav, A.; Muller, R.; Waters, K. M.; Shi, W.; Liu, X.; Zhang, L.; Knight, R.; Jensen, P. R.; Palsson, B. O.; Pogliano, K.; Linington, R. G.; Gutierrez, M.; Lopes, N. P.; Gerwick, W. H.; Moore, B. S.; Dorrestein, P. C.; Bandeira, N., Sharing and community curation of mass spectrometry data with Global Natural Products Social Molecular Networking. Nat Biotechnol 2016, 34, (8), 828–837.

39. Bender, D. A., Biochemistry of tryptophan in health and disease. Mol Aspects Med 1983, 6, (2), 101–97.

40. Poyatos, J. F.; Hurst, L. D., How biologically relevant are interaction-based modules in protein networks? Genome Biol 2004, 5, (11), R93.

41. Muller, A. M.; Hermanns, M. I.; Skrzynski, C.; Nesslinger, M.; Muller, K. M.; Kirkpatrick, C. J., Expression of the endothelial markers PECAM-1, vWf, and CD34 in vivo and in vitro. Exp Mol Pathol 2002, 72, (3), 221–9.

42. Aird, W. C., Phenotypic heterogeneity of the endothelium: II. Representative vascular beds. Circ Res 2007, 100, (2), 174–90.

43. Aird, W. C., Endothelial cell heterogeneity. Cold Spring Harb Perspect Med 2012, 2, (1), a006429.

44. Widlansky, M. E.; Gokce, N.; Keaney, J. F., Jr.; Vita, J. A., The clinical implications of endothelial dysfunction. J Am Coll Cardiol 2003, 42, (7), 1149–60.

45. Ganz, P.; Vita, J. A., Testing endothelial vasomotor function: nitric oxide, a multipotent molecule. Circulation 2003, 108, (17), 2049–53.

46. Paulus, W. J.; Vantrimpont, P. J.; Shah, A. M., Paracrine coronary endothelial control of left ventricular function in humans. Circulation 1995, 92, (8), 2119–26.

47. Abe, M.; Ono, J.; Sato, Y.; Okeda, T.; Takaki, R., Effects of glucose and insulin on cultured human microvascular endothelial cells. Diabetes Res Clin Pract 1990, 9, (3), 287–95.

48. Du, X. L.; Sui, G. Z.; Stockklauser-Farber, K.; Weiss, J.; Zink, S.; Schwippert, B.; Wu, Q. X.; Tschope, D.; Rosen, P., Introduction of apoptosis by high proinsulin and glucose in cultured human umbilical vein endothelial cells is mediated by reactive oxygen species. Diabetologia 1998, 41, (3), 249–56.

49. Graier, W. F.; Grubenthal, I.; Dittrich, P.; Wascher, T. C.; Kostner, G. M., Intracellular mechanism of high D-glucose-induced modulation of vascular cell proliferation. Eur J Pharmacol 1995, 294, (1), 221–9.

50. Kamal, K.; Du, W.; Mills, I.; Sumpio, B. E., Antiproliferative effect of elevated glucose in human microvascular endothelial cells. J Cell Biochem 1998, 71, (4), 491–501.

51. Lorenzi, M.; Nordberg, J. A.; Toledo, S., High glucose prolongs cell-cycle traversal of cultured human endothelial cells. Diabetes 1987, 36, (11), 1261–7.

52. Quagliaro, L.; Piconi, L.; Assaloni, R.; Martinelli, L.; Motz, E.; Ceriello, A., Intermittent high glucose enhances apoptosis related to oxidative stress in human umbilical vein endothelial cells: the role of protein kinase C and NAD(P)H-oxidase activation. Diabetes 2003, 52, (11), 2795–804.

53. McGinn, S.; Poronnik, P.; King, M.; Gallery, E. D.; Pollock, C. A., High glucose and endothelial cell growth: novel effects independent of autocrine TGF-beta 1 and hyperosmolarity. Am J Physiol Cell Physiol 2003, 284, (6), C1374–86.

54. Yuan, W.; Zhang, J.; Li, S.; Edwards, J. L., Amine metabolomics of hyperglycemic endothelial cells using capillary LC-MS with isobaric tagging. J Proteome Res 2011, 10, (11), 5242–50.

55. Chen, S.; Akter, S.; Kuwahara, K.; Matsushita, Y.; Nakagawa, T.; Konishi, M.; Honda, T.; Yamamoto, S.; Hayashi, T.; Noda, M.; Mizoue, T., Serum amino acid profiles and risk of type 2 diabetes among Japanese adults in the Hitachi Health Study. Sci Rep 2019, 9, (1), 7010.

56. Lai, M.; Liu, Y.; Ronnett, G. V.; Wu, A.; Cox, B. J.; Dai, F. F.; Rost, H. L.; Gunderson, E. P.; Wheeler, M. B., Amino acid and lipid metabolism in post-gestational diabetes and progression to type 2 diabetes: A metabolic profiling study. PLoS Med 2020, 17, (5), e1003112.

57. Lu, Y.; Wang, Y.; Liang, X.; Zou, L.; Ong, C. N.; Yuan, J. M.; Koh, W. P.; Pan, A., Serum Amino Acids in Association with Prevalent and Incident Type 2 Diabetes in A Chinese Population. Metabolites 2019, 9, (1).

58. Menni, C.; Fauman, E.; Erte, I.; Perry, J. R.; Kastenmuller, G.; Shin, S. Y.; Petersen, A. K.; Hyde, C.; Psatha, M.; Ward, K. J.; Yuan, W.; Milburn, M.; Palmer, C. N.; Frayling, T. M.; Trimmer, J.; Bell, J. T.; Gieger, C.; Mohney, R. P.; Brosnan, M. J.; Suhre, K.; Soranzo, N.; Spector, T. D., Biomarkers for type 2 diabetes and impaired fasting glucose using a nontargeted metabolomics approach. Diabetes 2013, 62, (12), 4270–6.

59. Wang, T. J.; Larson, M. G.; Vasan, R. S.; Cheng, S.; Rhee, E. P.; McCabe, E.; Lewis, G. D.; Fox, C. S.; Jacques, P. F.; Fernandez, C.; O’Donnell, C. J.; Carr, S. A.; Mootha, V. K.; Florez, J. C.; Souza, A.; Melander, O.; Clish, C. B.; Gerszten, R. E., Metabolite profiles and the risk of developing diabetes. Nat Med 2011, 17, (4), 448–53.

60. Koziel, A.; Woyda-Ploszczyca, A.; Kicinska, A.; Jarmuszkiewicz, W., The influence of high glucose on the aerobic metabolism of endothelial EA.hy926 cells. Pflugers Arch 2012, 464, (6), 657–69.

61. Badawy, A. A., Kynurenine Pathway of Tryptophan Metabolism: Regulatory and Functional Aspects. Int J Tryptophan Res 2017, 10, 1178646917691938.

62. Pedersen, E. R.; Tuseth, N.; Eussen, S. J.; Ueland, P. M.; Strand, E.; Svingen, G. F.; Midttun, O.; Meyer, K.; Mellgren, G.; Ulvik, A.; Nordrehaug, J. E.; Nilsen, D. W.; Nygard, O., Associations of plasma kynurenines with risk of acute myocardial infarction in patients with stable angina pectoris. Arterioscler Thromb Vasc Biol 2015, 35, (2), 455–62.

63. Sulo, G.; Vollset, S. E.; Nygard, O.; Midttun, O.; Ueland, P. M.; Eussen, S. J.; Pedersen, E. R.; Tell, G. S., Neopterin and kynurenine-tryptophan ratio as predictors of coronary events in older adults, the Hordaland Health Study. Int J Cardiol 2013, 168, (2), 1435–40.

64. Polyzos, K. A.; Ketelhuth, D. F., The role of the kynurenine pathway of tryptophan metabolism in cardiovascular disease. An emerging field. Hamostaseologie 2015, 35, (2), 128–36.

65. Aquilano, K.; Baldelli, S.; Ciriolo, M. R., Glutathione: new roles in redox signaling for an old antioxidant. Front Pharmacol 2014, 5, 196.

66. Yuan, W.; Edwards, J. L., Thiol metabolomics of endothelial cells using capillary liquid chromatography mass spectrometry with isotope coded affinity tags. J Chromatogr A 2011, 1218, (18), 2561–8.

67. Weidig, P.; McMaster, D.; Bayraktutan, U., High glucose mediates pro-oxidant and antioxidant enzyme activities in coronary endothelial cells. Diabetes Obes Metab 2004, 6, (6), 432–41.

68. Felice, F.; Lucchesi, D.; di Stefano, R.; Barsotti, M. C.; Storti, E.; Penno, G.; Balbarini, A.; Del Prato, S.; Pucci, L., Oxidative stress in response to high glucose levels in endothelial cells and in endothelial progenitor cells: evidence for differential glutathione peroxidase-1 expression. Microvasc Res 2010, 80, (3), 332–8.

69. Kashiwagi, A.; Asahina, T.; Ikebuchi, M.; Tanaka, Y.; Takagi, Y.; Nishio, Y.; Kikkawa, R.; Shigeta, Y., Abnormal glutathione metabolism and increased cytotoxicity caused by H2O2 in human umbilical vein endothelial cells cultured in high glucose medium. Diabetologia 1994, 37, (3), 264–9.

70. Hanschmann, E. M.; Godoy, J. R.; Berndt, C.; Hudemann, C.; Lillig, C. H., Thioredoxins, glutaredoxins, and peroxiredoxins--molecular mechanisms and health significance: from cofactors to antioxidants to redox signaling. Antioxid Redox Signal 2013, 19, (13), 1539–605.

71. Scocchi, M.; Tossi, A.; Gennaro, R., Proline-rich antimicrobial peptides: converging to a non-lytic mechanism of action. Cell Mol Life Sci 2011, 68, (13), 2317–30.

72. Migliaccio, A.; Castoria, G.; de Falco, A.; Bilancio, A.; Giovannelli, P.; Di Donato, M.; Marino, I.; Yamaguchi, H.; Appella, E.; Auricchio, F., Polyproline and Tat transduction peptides in the study of the rapid actions of steroid receptors. Steroids 2012, 77, (10), 974–8.

73. Radicioni, G.; Stringaro, A.; Molinari, A.; Nocca, G.; Longhi, R.; Pirolli, D.; Scarano, E.; Iavarone, F.; Manconi, B.; Cabras, T.; Messana, I.; Castagnola, M.; Vitali, A., Characterization of the cell penetrating properties of a human salivary proline-rich peptide. Biochim Biophys Acta 2015, 1848, (11 Pt A), 2868–77.

74. Vanhoof, G.; Goossens, F.; De Meester, I.; Hendriks, D.; Scharpe, S., Proline motifs in peptides and their biological processing. FASEB J 1995, 9, (9), 736–44.

75. Colombo, S.; Melo, T.; Martinez-Lopez, M.; Carrasco, M. J.; Domingues, M. R.; Perez-Sala, D.; Domingues, P., Phospholipidome of endothelial cells shows a different adaptation response upon oxidative, glycative and lipoxidative stress. Sci Rep 2018, 8, (1), 12365.

76. De Keyzer, D.; Karabina, S. A.; Wei, W.; Geeraert, B.; Stengel, D.; Marsillach, J.; Camps, J.; Holvoet, P.; Ninio, E., Increased PAFAH and oxidized lipids are associated with inflammation and atherosclerosis in hypercholesterolemic pigs. Arterioscler Thromb Vasc Biol 2009, 29, (12), 2041–6.

77. Tselepis, A. D.; John Chapman, M., Inflammation, bioactive lipids and atherosclerosis: potential roles of a lipoprotein-associated phospholipase A2, platelet activating factor-acetylhydrolase. Atheroscler Suppl 2002, 3, (4), 57–68.

78. Wang, A.; Dennis, E. A., Mammalian lysophospholipases. Biochim Biophys Acta 1999, 1439, (1), 1–16.

79. Marco-Ramell, A.; Palau-Rodriguez, M.; Alay, A.; Tulipani, S.; Urpi-Sarda, M.; Sanchez-Pla, A.; Andres-Lacueva, C., Evaluation and comparison of bioinformatic tools for the enrichment analysis of metabolomics data. BMC Bioinformatics 2018, 19, (1), 1.

80. Zhou, X.; Liao, W. J.; Liao, J. M.; Liao, P.; Lu, H., Ribosomal proteins: functions beyond the ribosome. J Mol Cell Biol 2015, 7, (2), 92–104.

81. Goldberg, A. L., Protein degradation and protection against misfolded or damaged proteins. Nature 2003, 426, (6968), 895–9.

82. Vinals, F.; Pouyssegur, J., Confluence of vascular endothelial cells induces cell cycle exit by inhibiting p42/p44 mitogen-activated protein kinase activity. Mol Cell Biol 1999, 19, (4), 2763–72.

83. Yu, Y.; Moulton, K. S.; Khan, M. K.; Vineberg, S.; Boye, E.; Davis, V. M.; O’Donnell, P. E.; Bischoff, J.; Milstone, D. S., E-selectin is required for the antiangiogenic activity of endostatin. Proc Natl Acad Sci U S A 2004, 101, (21), 8005–10.

84. Brigstock, D. R., Regulation of angiogenesis and endothelial cell function by connective tissue growth factor (CTGF) and cysteine-rich 61 (CYR61). Angiogenesis 2002, 5, (3), 153–65.

85. Elmasri, H.; Ghelfi, E.; Yu, C. W.; Traphagen, S.; Cernadas, M.; Cao, H.; Shi, G. P.; Plutzky, J.; Sahin, M.; Hotamisligil, G.; Cataltepe, S., Endothelial cell-fatty acid binding protein 4 promotes angiogenesis: role of stem cell factor/c-kit pathway. Angiogenesis 2012, 15, (3), 457–68.

86. Quinn, M. T.; Schepetkin, I. A., Role of NADPH oxidase in formation and function of multinucleated giant cells. J Innate Immun 2009, 1, (6), 509–26.

87. Holt, D. J.; Grainger, D. W., Multinucleated giant cells from fibroblast cultures. Biomaterials 2011, 32, (16), 3977–87.

88. Tse, G. M.; Law, B. K.; Chan, K. F.; Mas, T. K., Multinucleated stromal giant cells in mammary phyllodes tumours. Pathology 2001, 33, (2), 153–6.

89. Celton-Morizur, S.; Merlen, G.; Couton, D.; Desdouets, C., Polyploidy and liver proliferation: central role of insulin signaling. Cell Cycle 2010, 9, (3), 460–6.

