## Supporting Information for "Comprehensive multi-omics study of the molecular perturbations induced by simulated diabetes on coronary artery endothelial cells"

**This PDF file includes:**

Supplemental Experimental Methods
Supplemental References

Supplemental Figures 1-2:

**Figure S1.** Proteomics data normalization results using NormalyzerDE. (A) Total intensity of raw data before normalization. (B) Quantitative parameters of normalization algorithms (pooled intragroup coefficient of variation [PCV], median absolute deviation [PMAD], estimate of variance [PEV]). Qualitative parameters of normalization algorithms; (C) Box plots (D) MA plots, and (E) Density plots.

**Figure S2. Cellular confluence in control and experimental group.** Representative micrographs of Bovine Coronary Artery Endothelial Cells (BCAEC) cultured for 9 days with 5.5 mmol/L glucose (control group) and 20 mmol/L glucose+100 nmol/L insulin (simulated diabetes or experimental group). Images were taken using an EVOS® FLoid® Cell Imaging Station with a fixed 20x air objective. Abbreviations: NG, normal glucose; HG, high glucose; HI, high insulin.

**Table S2.** List of putatively annotated ($MS^2$ spectral matching) metabolites modulated by simulated diabetes.

**Table S4** Putative annotated proline-peptides altered by simulated diabetes in Bovine Coronary Artery Endothelial Cells by ProteinPilot Software and manual inspection.

In addition, this manuscript also has associated 3 .xlsx files corresponding to **Table S1, S3, and S5**.

**Table S1.** List of all the putatively annotated metabolites by $MS^2$ spectral matching against GNPS public spectral libraries.

**Table S3.** List of all detected peptides by ProteinPilot Software using the metabolomics datasets.

**Table S5.** List of the detected peptides and proteins in all conditions for SWATH-based quantification.

### SUPPLEMENTAL EXPERIMENTAL METHODS

**Metabolomics data processing.** For the XCMS pipeline, raw LC-$MS^2$ data files (.wiff and .wiff scan) were uploaded to the XCMS web-based platform at https://xcmsonline.scripps.edu (1). Preset parameters for TripleTOF 5600 in positive ionization mode and pairwise (unpaired t test) were selected, with the following modifications; min peak width, 15 s; max peak width, 90 s; feature detection mass tolerance, 15 ppm; and peak normalization, median fold change. For the MZmine pipeline, raw data files (.wiff and .wiff scan format) were first converted to .mzML using MSconvert (ProteoWizard) (2). Mass detection, chromatogram building and deconvolution (ADAP algorithm), isotopic assignment, feature alignment and gap-filling (to detect features missed during the initial alignment) were performed in MZmine Version 2.38 (3). Features with at least 2 isotopes were kept and the list containing all $MS^1$ was exported as a .csv file. The list comprising all the features detected by MZmine was used for comparison against the XCMS pipeline. For subsequent analysis, features detected in mobile phase (water/acetonitrile 95:5 v/v with 0.1% formic) blanks were considered contaminants and removed from samples using the *peak list row filter* tool within MZmine and manual inspection. Features with $MS^2$ data were kept. The peak areas and $MS^2$ data of filtered features were exported as .csv and .mgf files, respectively. This was followed by Feature Based Molecular Networking (FBMN) and spectral library matching performed into the Global Natural Products Social Molecular Networking (GNPS) web-platform (www.gnps.ucsd.edu) (4). The parameters used within the GNPS were precursor ion and product ion mass tolerance of 0.02 Da. The molecular network was created using a minimum cosine score of 0.6 and a minimum of 4 peaks matched. The spectra in the network were searched against GNPS spectral libraries using a minimum cosine score of 0.6 and at least 4 matched peaks as filters. To expand the annotation of the metabolites not automatically retrieved by spectral matching, the GNPS *in silico* tool, Network Annotation Propagation (NAP), was used (5). NAP uses the output of molecular networking to rerank *in silico* candidate structure lists, which has been shown to improve metabolite annotation. For NAP, adducts $[M+NH_4]+$, [M+H]+, [M+Na]+, and [M+K]+ with *m/z* tolerance set to 10 ppm were searched using the parameters described at: https://proteomics2.ucsd.edu/ProteoSAFe/status.jsp?task=96cda48c0df64d3398a8f9088907afb5. A chemical class was automatically assigned to all putative annotations obtained through GNPS library matching and NAP, using the Classyfire chemical ontology nomenclature (6). Molecular networking, NAP, and Classyfire outputs were integrated using the MolNetEnhancer workflow to visualize the metabolome detected (7). As a complementary step to annotate metabolites, chemical substructures were recognized using the MS2LDA computational tool aiming to discover co-occurring fragments and neutral losses (referred as to Mass2Motifs [M2M]) within the MS2 data (using the same .mgf file as in GNPS). This provides chemical information at the substructure level of the metabolites (8). $MS^1$ quantification (with normalization “on”) was utilized to visualize relevant sub structurally-related features in MS2LDA. Fragments retrieved by M2M of interest were searched against the mzCloud database (https://www.mzcloud.org) to obtain Heuristic and Quantum Chemical predictions (non-trivial *in silico* predictions).

To compare the features detected by both platforms, the lists of $MS^1$ were transformed to mzTab format and imported into MZmine for feature alignment using a tolerance of 0.01 m/z and 1 min. The aligned features were considered the same entity. A Venn diagram was used to visualize and compare the features detected by MZmine and XCMS software.

### REFERENCES


1. Gowda, H.; Ivanisevic, J.; Johnson, C. H.; Kurczy, M. E.; Benton, H. P.; Rinehart, D.; Nguyen, T.; Ray, J.; Kuehl, J.; Arevalo, B.; Westenskow, P. D.; Wang, J.; Arkin, A. P.; Deutschbauer, A. M.; Patti, G. J.; Siuzdak, G., Interactive XCMS Online: simplifying advanced metabolomic data processing and subsequent statistical analyses. *Anal Chem* **2014,** 86, (14), 6931-9.
2. Holman, J. D.; Tabb, D. L.; Mallick, P., Employing ProteoWizard to Convert Raw Mass Spectrometry Data. *Curr Protoc Bioinformatics* **2014,** 46, 13 24 1-9.
3. Pluskal, T.; Castillo, S.; Villar-Briones, A.; Oresic, M., MZmine 2: modular framework for processing, visualizing, and analyzing mass spectrometry-based molecular profile data. *BMC Bioinformatics* **2010,** 11, 395.
4. Nothias, L. F.; Petras, D.; Schmid, R.; Duhrkop, K.; Rainer, J.; Sarvepalli, A.; Protsyuk, I.; Ernst, M.; Tsugawa, H.; Fleischauer, M.; Aicheler, F.; Aksenov, A. A.; Alka, O.; Allard, P. M.; Barsch, A.; Cachet, X.; Caraballo-Rodriguez, A. M.; Da Silva, R. R.; Dang, T.; Garg, N.; Gauglitz, J. M.; Gurevich, A.; Isaac, G.; Jarmusch, A. K.; Kamenik, Z.; Kang, K. B.; Kessler, N.; Koester, I.; Korf, A.; Le Gouellec, A.; Ludwig, M.; Martin, H. C.; McCall, L. I.; McSayles, J.; Meyer, S. W.; Mohimani, H.; Morsy, M.; Moyne, O.; Neumann, S.; Neuweger, H.; Nguyen, N. H.; Nothias-Esposito, M.; Paolini, J.; Phelan, V. V.; Pluskal, T.; Quinn, R. A.; Rogers, S.; Shrestha, B.; Tripathi, A.; van der Hooft, J. J. J.; Vargas, F.; Weldon, K. C.; Witting, M.; Yang, H.; Zhang, Z.; Zubeil, F.; Kohlbacher, O.; Bocker, S.; Alexandrov, T.; Bandeira, N.; Wang, M.; Dorrestein, P. C., Feature-based molecular networking in the GNPS analysis environment. *Nat Methods* **2020,** 17, (9), 905-908.
5. da Silva, R. R.; Wang, M.; Nothias, L. F.; van der Hooft, J. J. J.; Caraballo-Rodriguez, A. M.; Fox, E.; Balunas, M. J.; Klassen, J. L.; Lopes, N. P.; Dorrestein, P. C., Propagating annotations of molecular networks using in silico fragmentation. *PLoS Comput Biol* **2018,** 14, (4), e1006089.
6. Djoumbou Feunang, Y.; Eisner, R.; Knox, C.; Chepelev, L.; Hastings, J.; Owen, G.; Fahy, E.; Steinbeck, C.; Subramanian, S.; Bolton, E.; Greiner, R.; Wishart, D. S., ClassyFire:

automated chemical classification with a comprehensive, computable taxonomy. *J Cheminform* **2016,** 8, 61.
7. Ernst, M.; Kang, K. B.; Caraballo-Rodriguez, A. M.; Nothias, L. F.; Wandy, J.; Chen, C.; Wang, M.; Rogers, S.; Medema, M. H.; Dorrestein, P. C.; van der Hooft, J. J. J., MolNetEnhancer: Enhanced Molecular Networks by Integrating Metabolome Mining and Annotation Tools. *Metabolites* **2019,** 9, (7).
8. van der Hooft, J. J.; Wandy, J.; Barrett, M. P.; Burgess, K. E.; Rogers, S., Topic modeling for untargeted substructure exploration in metabolomics. *Proc Natl Acad Sci U S A* **2016,** 113, (48), 13738-13743.

### SUPPLEMENTAL FIGURES

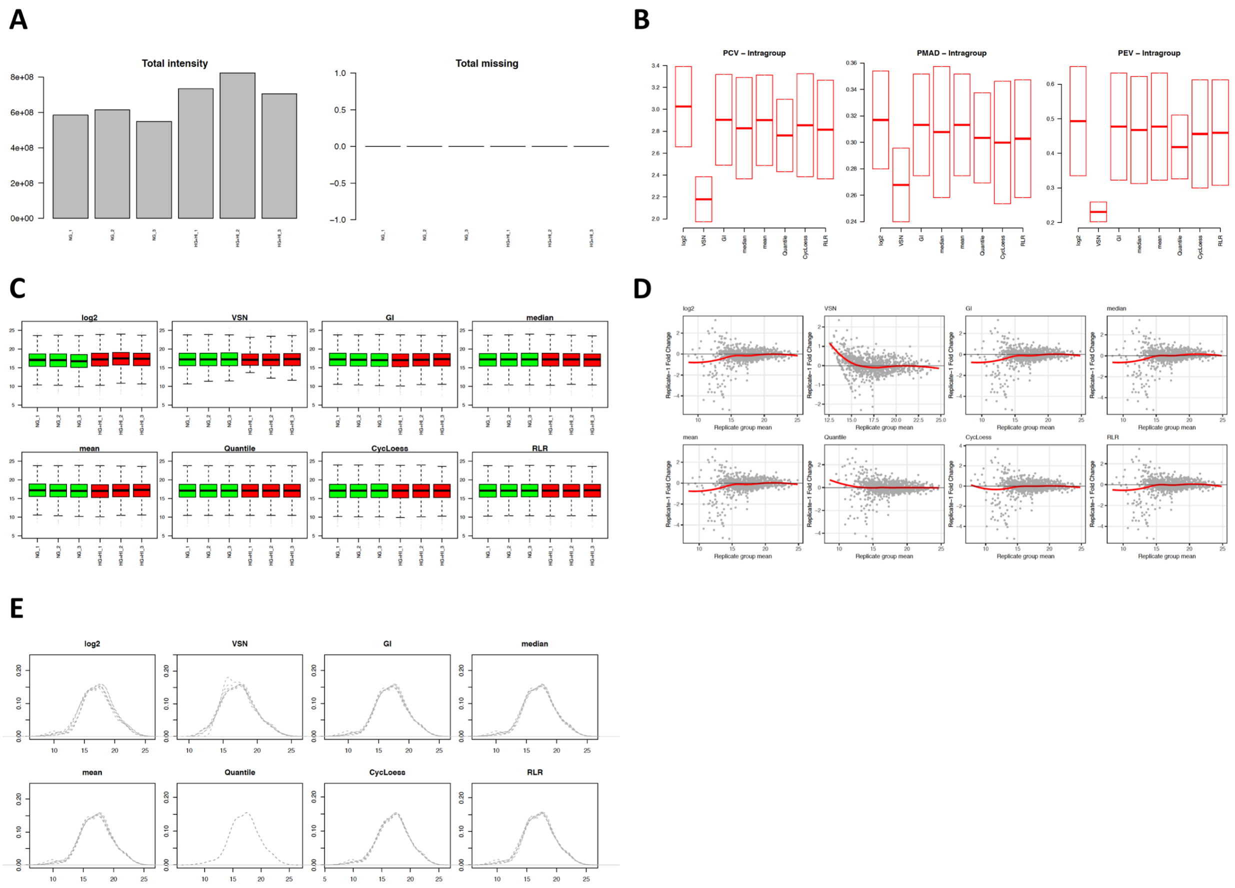


**Figure S1***.* Proteomics data normalization results using NormalyzerDE. (A) Total intensity of raw LC-MS[2] data before normalization. (B) Quantitative parameters of normalization algorithms (pooled intragroup coefficient of variation [PCV], median absolute deviation [PMAD], estimate of variance [PEV]). Qualitative parameters of normalization algorithms; (C) Box plots (D) MA plots, and (E) Density plots.

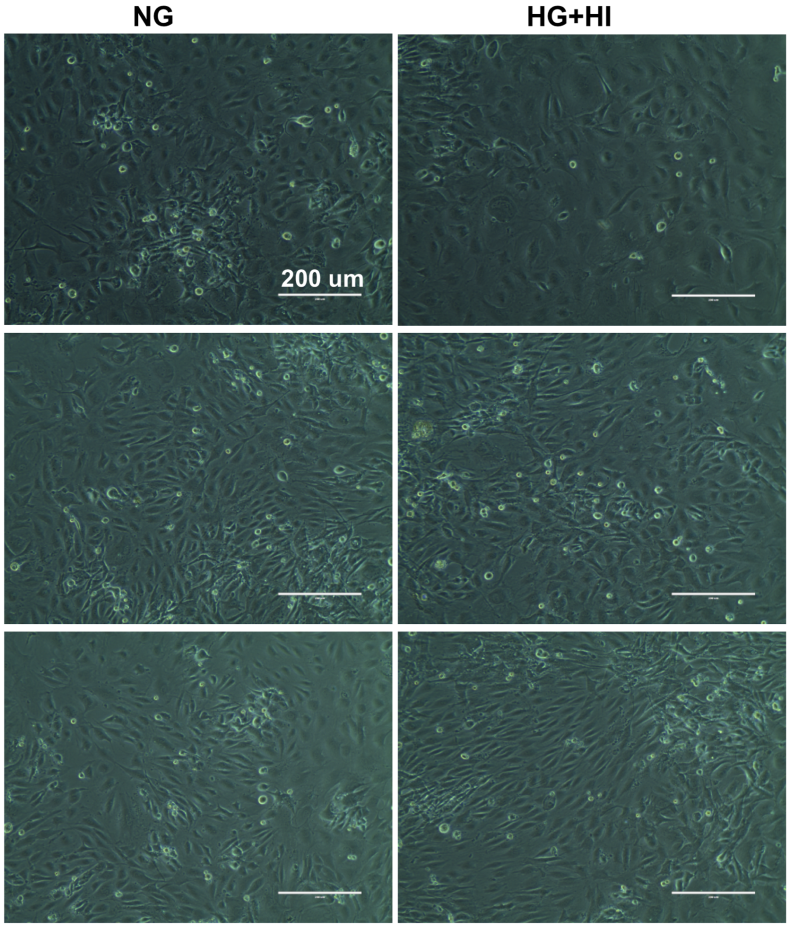


**Figure S2. Cellular confluence in control and experimental group.** Representative micrographs of Bovine Coronary Artery Endothelial Cells (BCAEC) cultured for 9 days with 5.5 mmol/L glucose (control group) and 20 mmol/L glucose+100 nmol/L insulin (simulated diabetes or experimental group). Images were taken using an EVOS® FLoid® Cell Imaging Station with a fixed 20x air objective. Abbreviations: NG, normal glucose; HG, high glucose; HI, high insulin.

### SUPPLEMENTAL TABLES

**Table S2. List of putatively annotated (MS2 spectral matching) metabolites modulated by simulated diabetes.**

| | **Compound name** | **Chemical subclass[a]** | **Retention time** | **Experimental mass [adduct]** | **Exact mass** | **Mass error (ppm)** | **Fold change (vs. control)[b]** |
|---|---|---|---|---|---|---|---|
| **Up-regulated** | Kynurenine | Carbonyl compounds | 3.2 | 209.0914 $[M+H]^+$ | 209.0926 | -5.7 | 4.2 |
| | Panthothenic acid | Alcohols and polyols | 2.8 | 242.0992 $[M+Na]^+$ | 242.1004 | -4.9 | 1.38 |
| | Threonine | Aminoacids, peptides and analogues | 0.6 | 120.0658 $[M+H]^+$ | 120.0661 | -2.4 | 1.4 |
| | Valine | Aminoacids, peptides and analogues | 0.75 | 118.086 $[M+H]^+$ | 118.0868 | -6.7 | 1.5 |
| | Proline | Aminoacids, peptides and analogues | 0.6 | 116.0710 $[M+H]^+$ | 116.0712 | -1.7 | 1.7 |
| | Leucine | Aminoacids, peptides and analogues | 1.3 | 132.1022 $[M+H]^+$ | 132.1025 | -2.2 | 1.37 |
| | Serine | Aminoacids, peptides and analogues | 0.6 | 106.0504 $[M+H]^+$ | 106.0504 | 0 | 1.5 |
| | Glutamic acid | Aminoacids, peptides and analogues | 0.6 | 148.0604 $[M+H]^+$ | 148.061 | -4.0 | 1.6 |
| | Methionine | Aminoacids, peptides and analogues | 0.9 | 150.0584 $[M+H]^+$ | 150.0589 | -3.3 | 1.4 |
| | Tyrosine | Aminoacids, peptides and analogues | 1.3 | 182.0810 $[M+H]^+$ | 182.0817 | -3.8 | 1.3 |
| | Glutathione | Aminoacids, peptides and analogues | 1.3 | 615.1719 $[2M+H]^+$ | 615.1755 | -5.8 | 2.1 |

| | | | | | | | |
|---|---|---|---|---|---|---|---|
| | Glutamyl-phenylalanine | Aminoacids, peptides and analogues | 9.4 | 295.1285 $[M+H]^+$ | 295.1294 | -3.0 | 1.4 |
| | 2-aminoadipate | Aminoacids, peptides and analogues | 0.6 | 162.076 $[M+H]^+$ | 162.0767 | -4.3 | 1.4 |
| | 2,4-dimethylbenzoic acid | Benzoic acids and derivatives | 15.4 | 151.0753 $[M+H]^+$ | 151.0759 | -3.9 | 1.4 |
| | Phosphocholine | Quaternary ammonium salts | 0.75 | 184.0730 $[M]^+$ | 184.0738 | -4.3 | 1.8 |
| **Down-regulated** | | | | | | | |
| | PC(18:1(9Z)/18:1(9Z))[d] | Glycerophosphocholines | 24.4 | 786.5972 $[M+H]^+$ | 786.6013 | -5.2 | -5.8 |
| | PC(16:0/18:1(9Z))[d] | Glycerophosphocholines | 24.8 | 760.5816 $[M+H]^+$ | 760.5857 | -5.3 | -21.6 |
| | (2S)-2-(6-Hydroxy-6-methyloctyl)-2H-furan-5-one | Furanones | 15.4 | 244.1906 $[M+NH_4]^+$ | 244.1912 | -2.4 | -1.86 |

[a] Classification by Classyfire http://classyfire.wishartlab.com
[b] Fold change of HG+HI/NG
[c] Structure annotated by spectral match against GNPS public spectral libraries and validated by manual inspection of MS2 data
[d] The exact position and the cis/trans configuration could not be determined.

**Table S4. Putative annotated proline-peptides altered by simulated diabetes in Bovine Coronary Artery Endothelial Cells by ProteinPilot Software and manual inspection**

| Sequence | Gene[a] | Experimental mass [adduct] | Exact mass | Mass error (ppm) | Retention Time (min) | Fold change (vs. control)[b] | p-value | Confidence (%)[c] |
|---|---|---|---|---|---|---|---|---|
| TAPEIAVP | UFC1 | 399.2232 $[M+2H]^+$ | 399.2238 $[M+2H]^+$ | -1.5 | 9.4 | 1.46 | 0.04 | 99 |
| PPPPVP[ox]PPPPPP | WASF1 | 601.3339 $[M+2H]^+$ | 601.3344 $[M+2H]^+$ | -0.8 | 9.9 | 1.27 | 0.03 | 94.7 |
| LPP | Unknown | 326.2067 $[M+H]^+$ | 326.2074 $[M+H]^+$ | -2.2 | 8.4 | 1.45 | 0.01 | Manual inspection |

[ox]*oxidized residue*

[a] Gene ID of the corresponding protein contained the peptide sequence
[b]*high-glucose+high insulin*$_{AUC}$*/normal glucose*$_{AUC}$
[c]*Confidence threshold by ProteinPilot algorithm*
